# Protein recruitment to dynamic DNA-RNA host condensates

**DOI:** 10.1101/2024.06.04.597281

**Authors:** Mahdi Dizani, Daniela Sorrentino, Siddharth Agarwal, Jaimie Marie Stewart, Elisa Franco

## Abstract

We describe the design and characterization of artificial nucleic acid condensates that are engineered to recruit and locally concentrate proteins of interest *in vitro*. These condensates emerge from the programmed interactions of nanostructured motifs assembling from three DNA strands and one RNA strand that can include an aptamer domain for the recruitment of a target protein. Because condensates are designed to form regardless of the presence of target protein, they function as “host” compartments. As a model protein we consider streptavidin (SA) due to its widespread use in binding assays, thus the host condensates presented here could find immediate use for the physical separation of a variety of biotin-tagged components. In addition to demonstrating protein recruitment, we describe two approaches to control the onset of condensation and protein recruitment. The first approach uses UV irradiation, a physical stimulus that bypasses the need for exchanging molecular inputs and is particularly convenient to control condensation in emulsion droplets. The second approach uses RNA transcription, a ubiquitous biochemical reaction that is central to the development of the next generation of living materials. We finally show that the combination of RNA transcription and degradation leads to an autonomous dissipative system in which host condensates and protein recruitment occur transiently, and that the host condensate size as well as the timescale of the transient can be controlled by the level of RNA degrading enzyme.

**For Table of Contents Only:** 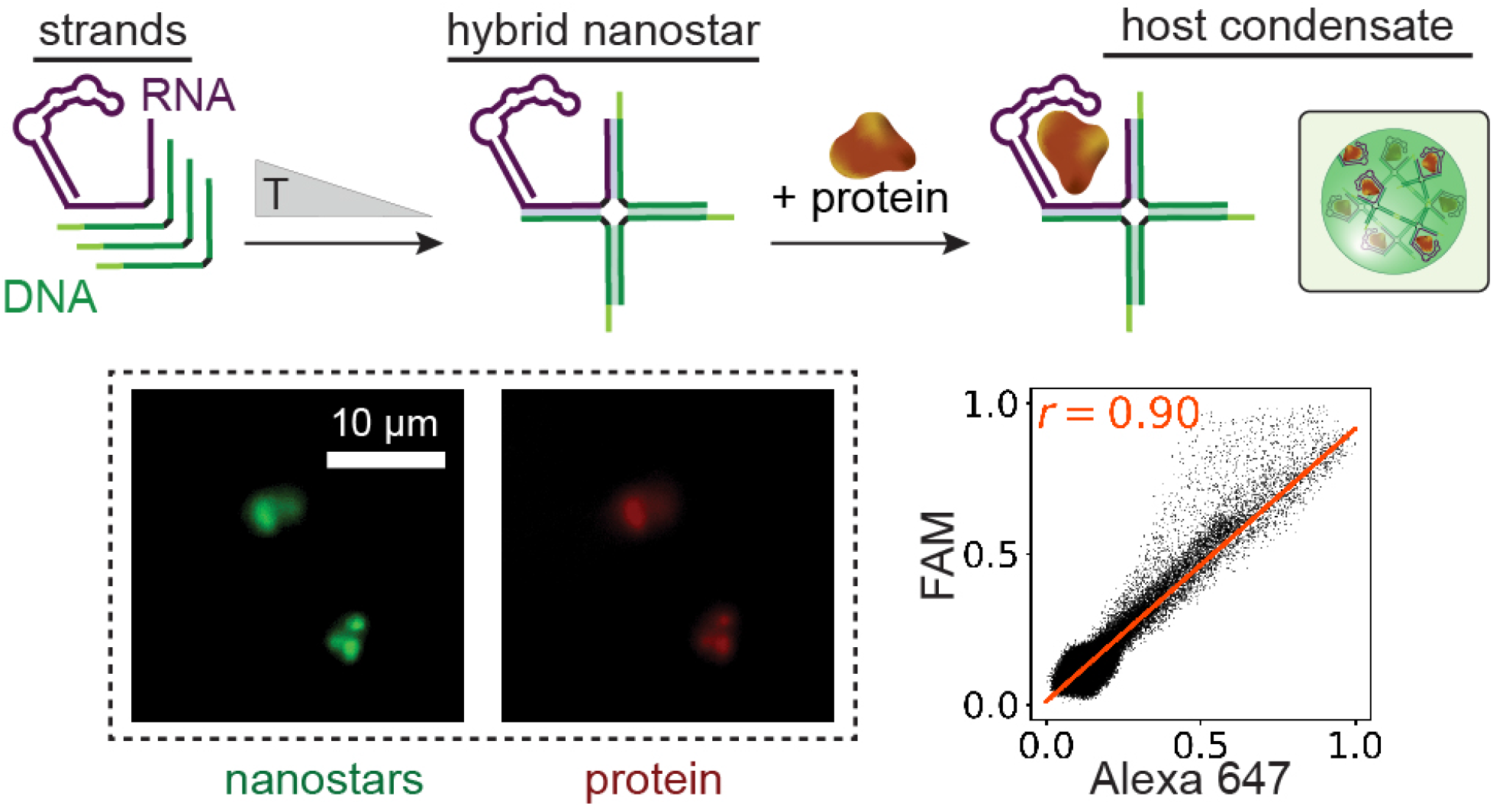

## Introduction

The ability to separate, sort and spatially organize molecules is central to biological manufacturing and synthetic biology. Of particular interest is the development of low-cost, low-volume bioreactors forming on-demand to handle scarce, toxic, or high-value molecules^1, 2^. In this context, methods relying on spontaneous biochemical separation, rather than physical separation, are highly sought after. Biological cells have mastered this task by evolving machinery to autonomously segregate DNA, proteins, small molecules, and entire reaction pathways through phase separation^3^. Droplet-like organelles without a membrane, such as glycolytic bodies, stress granules, and nucleoli, spontaneously condense within cells, and recruit specific molecules and reaction pathways at distinct times and locations^4,5^. These organelles, also known as biomolecular condensates, can effectively “host” many molecular “guests” or “clients”, which do not contribute to the condensation process^4^. Taking inspiration from nature, artificial host condensates arising from the interactions of synthetic polymers, engineered proteins, nucleic acids, as well as mixtures of these components, have been developed to biochemically recruit and localize molecules and pathways^6–9^.

DNA and RNA are particularly versatile polymers to build artificial condensates^10,11^. Through sequence design, nucleic acid molecules can be optimized to have a target secondary structure as well as specific affinity for distinct molecules, permitting the synthesis of complexes that phase separate predictably. A productive approach to build DNA condensates is to use artificial motifs that include multiple strands of DNA that self-assemble into “nanostars”^12^. DNA nanostars interact via single-stranded sticky end overhangs at the tip of each arm, generating DNA dense compartments with variable viscoelastic properties that depend on the number of arms (typically 3-6), the arm length, and the sticky-end length and sequence^10^. The resulting DNA condensates can form and dissolve in response to changes in solvent and temperature^13^, to specific molecular inputs^14,15^, and to physical stimuli such as light^16,17^. Nanostars can be modified to include recruitment domains for other oligonucleotides, making it possible to incorporate strand displacement reactions^15^, and even RNA transcription^15,18^. RNA nanostars have been demonstrated to recruit specific proteins through aptamers^19, 20^. Thus, nucleic acid nanostars are an excellent approach to build host condensates for separation of biomolecules.

Here we describe a strategy to localize proteins to DNA host condensates through RNA aptamers (Figure 1). We modified DNA nanostars to include an RNA strand encoding an aptamer domain for the recruitment of streptavidin (SA), which is widely used in combination with biotin for a multitude of affinity purification assays^21^. We first characterize how the presence of the aptamer and recruitment of SA affect the growth rate of DNA condensates. We then demonstrate how DNA host condensate growth, and thus SA localization to the condensates, can be induced on-demand with physical or biomolecular stimuli. As a physical input, we use UV light to trigger host condensate formation by a modified DNA nanostar that includes a UV-cleavable domain. We also show that UV light is an expedient tool to trigger host condensates in water-in-oil emulsion droplets. As a biomolecular stimulus, we use *in-situ* transcription of the RNA strand needed to establish necessary DNA nanostar hybridization as well as recruitment of SA. We further demonstrate that in the presence of RNase H, host condensates form transiently through a dissipative process, in which condensate size and timescale of condensate presence can be controlled by changing the amount of RNase H.

**Figure 1.**
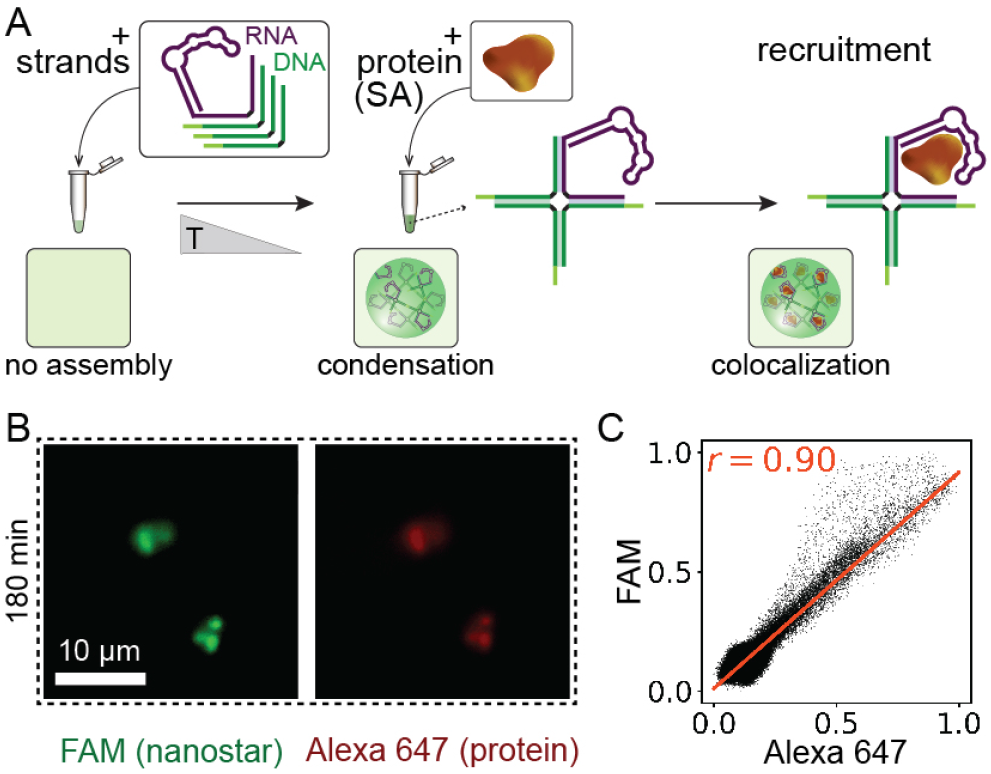
Selective protein recruitment via hybrid DNA-RNA nanostar. A) Schematic of 4-arm hybrid DNA-RNA nanostars that phase separate into protein-recruiting condensates. Our design comprises three DNA strands (green) modified at one end with single-stranded overhangs (sticky ends, light green) that allow condensates to assemble, and an RNA strand (purple) extended with an aptamer sequence responsible for protein recruitment (SA, brown). B) Representative fluorescence microscopy images showing the assembly of the hybrid DNA-RNA condensates (green, FAM) and the SA recruitment (red, Alexa 647) 180 minutes after the annealing step and isothermal addition of SA. C) Scatter plot of measured pixel intensity showing the correlation (Pearson’s correlation coefficient, r) of FAM (nanostar) and Alexa 647 (protein) fluorophores in a sample containing hybrid DNA-RNA condensates recruiting their corresponding client protein (SA). This correlation confirms that nanostars and proteins are colocalized. We used equimolar amounts of nanostar and protein, 2.5 μM.

Our approach is modular: the SA aptamer considered here could be replaced by aptamers recruiting other types of proteins^19,20^. Our host condensates form robustly under *in vitro* transcription conditions, and could be therefore used for synthetic biology applications to build dynamically controllable membraneless organelles for the localization of enzymes and reactions responding to transcriptional signals^22^.

## Results and discussion

### Hybrid DNA-RNA condensates can be programmed to recruit client proteins

We designed 4-arm hybrid DNA-RNA nanostars that interact via palindromic sticky ends to form condensates, and that through an encoded RNA aptamer can recruit streptavidin (SA) as an example target protein. The nanostar includes three DNA strands and one RNA strand which are partially complementary to one another and hybridize to form four arms, each 16 nucleotide (nt) long (Figure 1A). Palindromic sticky ends are 4-nt long (Figure 1A, light green domains) and are present at the 5’ end of the three DNA strands only (Figure 1A, green). We extended the 5’ end of the RNA strand to include an SA-binding aptamer^23^. Nanostars were assembled by thermal annealing of all the strands (each at 2.5 µM concentration) from 72 to 25 °C (at −1°C/min) in a buffer including 20 mM Tris-HCl and 12.5 mM MgCl_2_. Strand interactions and nanostar folding were verified through native PAGE, shown in the Supporting Information (SI) Figure S1. To quantify condensate formation, DNA-RNA nanostars were tagged with a fluorescently labeled DNA strand (0.5 µM of the total concentration of D_1_ is FAM-labeled), and SA was tagged with an orthogonal fluorophore (Alexa 647). This enabled condensate tracking via fluorescence microscopy (Figure 1B) as well as measuring DNA and protein colocalization by computing the Pearson’s correlation coefficient (r), which measures the strength of the linear relationship between the fluorescence intensity values of the green and red channels on the hybrid DNA-RNA condensates (Figure 1C). An r close to 0 indicates low colocalization of the fluorophores on the condensates, while a value close to 1 indicates high colocalization.

### Growth dynamics of DNA−RNA host condensates

We measured condensate formation and growth, as well as colocalization with SA, by tracking the samples over time via fluorescence microscopy experiments as shown in Figure 2. First, we measured how the presence of the RNA strand, the RNA aptamer, and the SA-aptamer complex affect condensate growth, when compared to all-DNA nanostars previously considered in the literature^24^. We tested different design variants that include the same 3 “core” strands D_1_, D_2_, D_3_ while the fourth strand is either: **(A)** a DNA sequence including the sticky end (D_4_, Figure 2A), **(B)** a blunt-ended DNA strand, without sticky end (D_4_^b^, Figure 2B), **(C)** a blunt-ended RNA strand (R_4_^b^, Figure 2C), **(D)** an RNA strand modified to contain the aptamer site for protein binding (R_4_^apt^, Figure 2D). Finally we considered the case in which the RNA aptamer and its target SA are both present (R_4_^apt^ + SA, Figure 2E), where SA was added immediately after annealing. RNA strands R_4_^b^ and R_4_^apt^ were purified with a column-based protocol. We imaged condensates within 15, 60 and 180 minutes after annealing completion, and we measured their area on the focal plane and number (Figure 2A-E).

**Figure 2.**
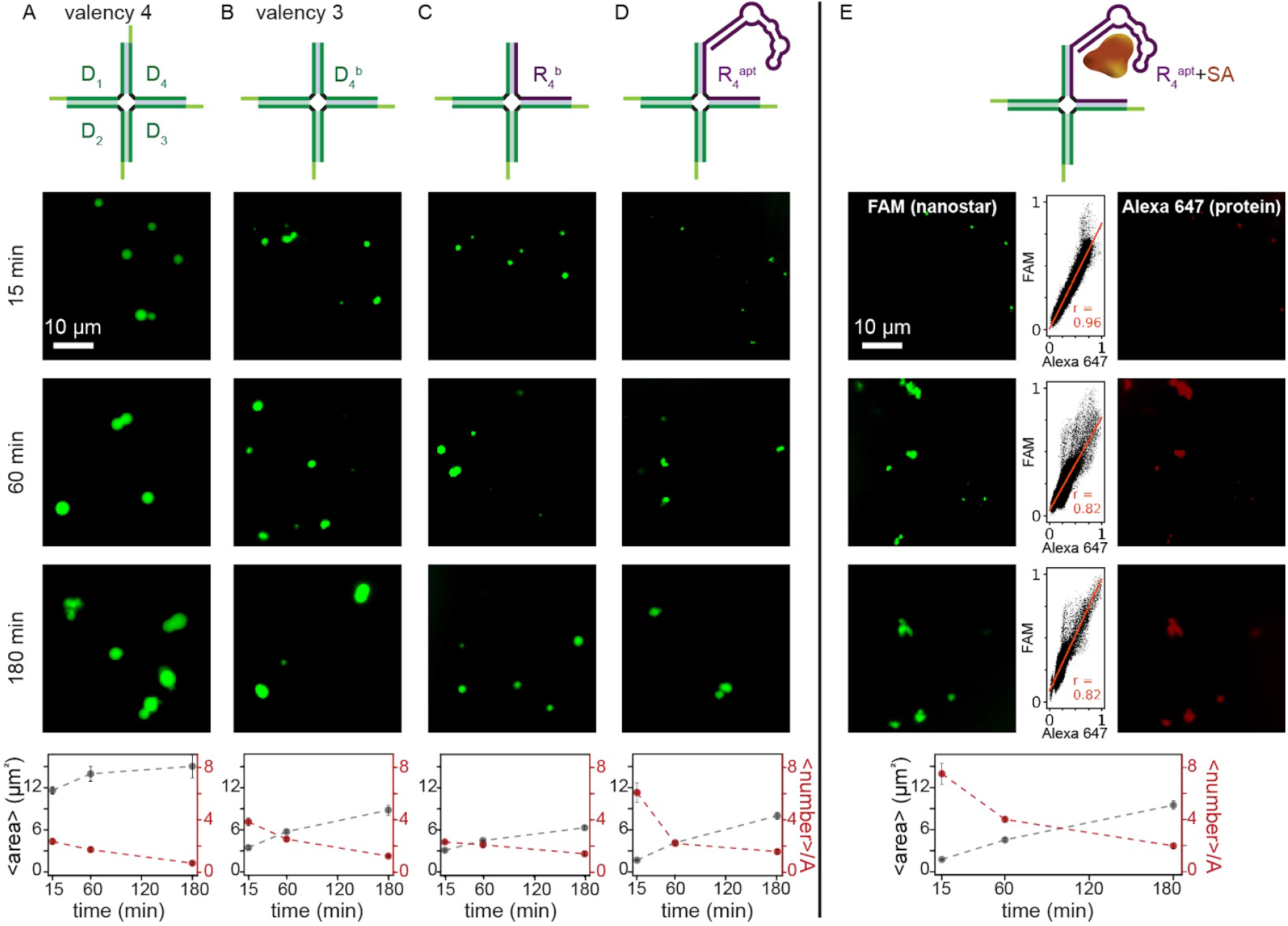
Growth of host condensates formed by DNA-RNA nanostars. We programmed nanostar-forming condensates using a ‘core structure’ (D_1_, D_2_, and D_3_, green) in which one strand can be replaced with an aptamer-carrying RNA strand. These experiments compare the growth rate of condensates emerging when using nanostar variants that differ in the nature and length of the fourth strand: A) a DNA sequence with sticky end (D_4_); B) a blunt DNA strand without sticky end (D_4_^b^); C) a blunt-ended RNA strand (R_4_^b^); D) an RNA strand modified to contain the aptamer site for protein binding (R_4_^apt^); E) an RNA-aptamer strand targeting the SA protein (R_4_^apt^ + SA). Plots at the bottom of each panel show average area and number density when incubated at room temperature (25 °C) for 180 minutes after annealing at strand concentrations of 2.5 μM. Number density is computed as the number of condensates per field of view of each image reported here (see Methods for additional detail). The numbers on the left of the images indicate the time (in minutes) at which these images were captured from the start of incubation (25 °C) after annealing completion. Dots correspond to the average of three experimental replicas (n = 3); error bars are the standard deviation of the mean.

Variants differ in terms of valency (the all-DNA design has valency 4, the others have valency 3), as well as in terms of the size and composition of the fourth arm, thus we expected different rates of condensate growth. All samples produced condensates under our experimental conditions (Figure 2A-E), confirming that three sticky ends (valency 3) are sufficient for condensation, and that nanostar interactions are not compromised by the presence of RNA, RNA aptamer, or SA. We found design (A) with four sticky ends (valency 4) to produce the fastest growing condensates, leading to the largest average area after 180 minutes of incubation (Figure 2A-E, bottom). At the same time, design (A) shows the lowest condensate number-density (computed as discussed in the Methods) which indicates that condensates fuse faster than other designs. Valency 3 DNA design (B), and hybrid DNA-RNA designs (C,D), yield condensates with a comparable average area near 4 µm^2^ after 180 minutes. The decrease in number density over time is again indicative of fusion events; the presence of SA RNA aptamer (with or without SA) appears to induce the emergence of a large number of small condensates, which grow and fuse more slowly (Figure 2D-E). Box plots of condensate area as well as total condensate numbers are in Figure S2.

Importantly, these experiments show that design (D), which includes the SA aptamer (strand R_4_^apt^), in the presence of SA produces condensates that recruit SA to the DNA dense phase, as shown by the representative images and the colocalization plots in Figure 2E. These condensates form robustly, and they grow at a speed comparable to the case in which SA is absent, indicating that our DNA-RNA nanostars are a viable motif to localize proteins to artificial membraneless compartments.

### UV light activation of nanostar condensation and protein recruitment

We next demonstrate that the timing of host condensate formation as well as client recruitment can be controlled by building nanostars whose interactions are regulated by an external input. We first consider UV light as a physical input, which is convenient as a “remote” control that does not require changes in the sample composition. Building on previous work^16^, we modified our nanostars design so that one of the three DNA arms (D_2_^photo^, Figure 3A) includes a UV-activatable sticky-end domain. This domain is designed to fold into a hairpin structure (red, Figure 3) preventing interactions with other sticky ends. This domain reduces the nanostar valency to 2, which is insufficient for condensation. The backbone of the hairpin stem contains a photocleavable (PC) linker (2-nitrobenzyl photolabile functional group) that is cleaved by UV light at a specific wavelength (300-350 nm)^25^. Upon UV-irradiation and cleavage of the protector strand, the hairpin unbinds from the sticky end bringing the nanostar valency to 3 and enabling the formation of condensates. Since the PC linker is unstable at high temperatures, the PC-modified strand was added isothermally at 25 °C after annealing^16^. First, we confirmed that in the absence of UV irradiation, no condensates form regardless of which RNA strand is included (Figure S3). Then we confirmed that UV irradiation of nanostar variants (C) or (D) (including respectively R_4_^b^ and R_4_^apt^) for 180 s triggers the formation of condensates in the absence or presence of SA (see Figure S4 for representative microscopy images). For design (C), the condensate average area increases over time, while the number density decreases indicating that fusion events occur (Figure 3B). Variant (D) that includes the SA aptamer shows slower initial growth, and the condensate number density increases first before decreasing due to fusion events (Figure S5 includes box plots of condensate size for each assay). When present in the sample, SA is recruited only to photoactivated condensates of variant (D) that include SA aptamer R_4_^apt^ resulting in significant colocalization (r=0.85 at 180 min). Colocalization analysis of 60 condensates in the absence and presence of SA is shown in Figure S6, confirming quantitatively that the aptamer is essential for protein recruitment. Overall, these results show that our photo-activatable nanostars partition the proteins into condensates only after UV irradiation, with growth kinetics comparable to the initial variants (Figure 2).

**Figure 3.**
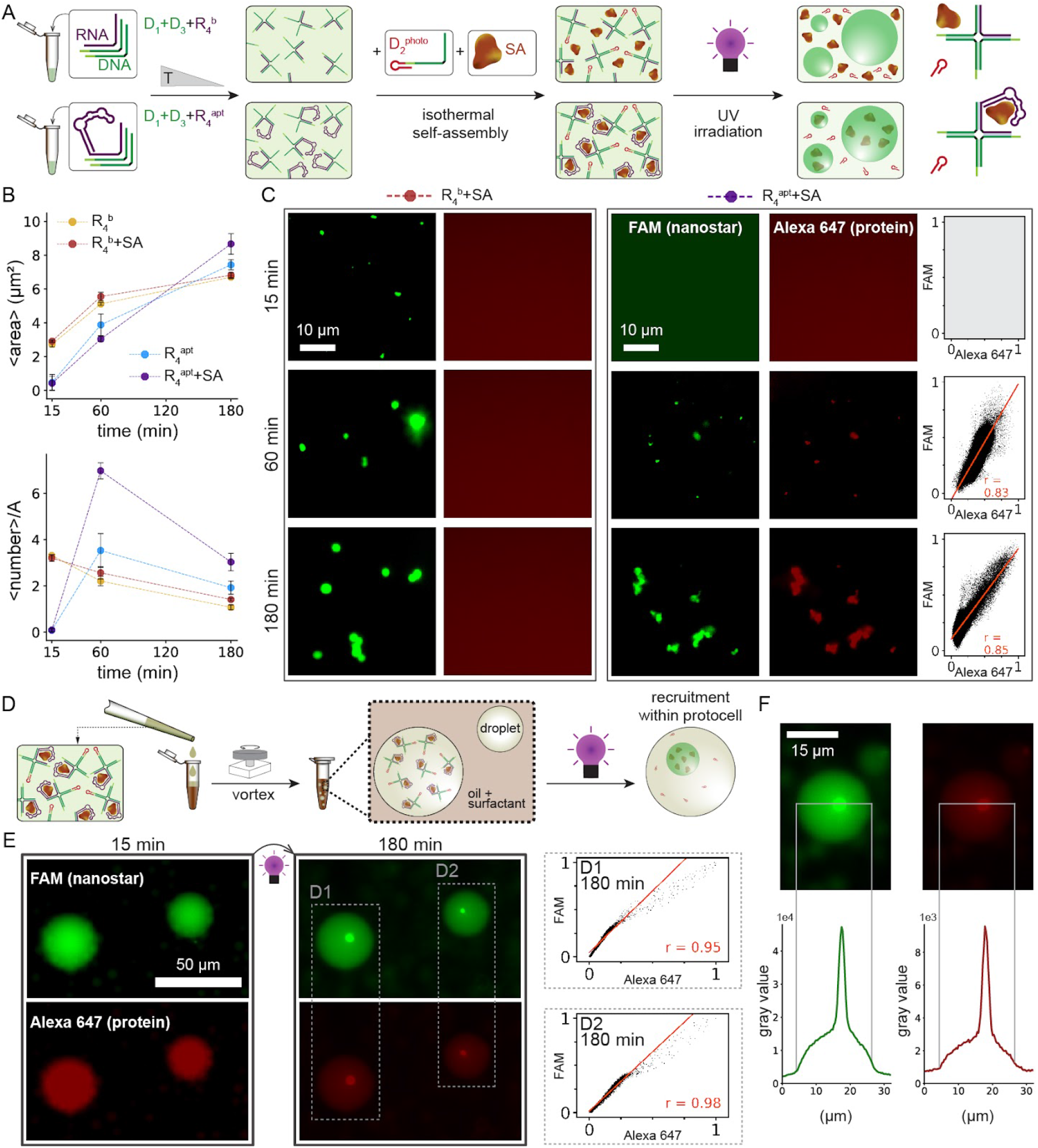
UV-controlled host condensate formation and protein capture. A) We modified strand D_2_ to include a hairpin domain that protects and inactivates the sticky end. The hairpin stem includes a UV-photocleavable linker: upon UV irradiation the protective domain unbinds from the sticky end changing nanostar valency from two to three and allowing the nanostar to form condensates. B) Plots of the average area and the number density of condensates up to 180 minutes after UV irradiation (irradiation time t_UV_ = 180 s) in the absence or presence of protein. Dots correspond to the average of three experimental replicas (n = 3); error bars are the standard deviation of the mean. C) Representative fluorescence microscopy images for activated nanostars phase separating into condensates. Protein capture is achieved only when the RNA strand contains an aptamer. Images were captured at 15, 60, and 180 minutes after UV irradiation at 25 °C. D) Schematic of the encapsulation process of hybrid DNA-RNA nanostars, as well as proteins, followed by protein recruitment within emulsion droplets. E) Example fluorescence images of hybrid DNA-RNA nanostars and their client protein inside droplets shown at 15 and 180 minutes after irradiation; correlation plots confirm that the reporters are colocalized. F) Intensity analysis of an example droplet 180 minutes after UV irradiation. The plots show the fluorescence intensity variation across an individual droplet, and clearly the peak intensity increases in both channels at the location of condensate.

A useful application of light-triggerable host condensates is that they could be activated on-demand in microcompartments that do not permit material exchange with the surrounding environment. Water-in-oil droplets are a versatile platform for high-throughput screening of reactions as well as minimalistic synthetic cells^26–29^. Building on our previous results, we demonstrate that nanostars encapsulated in water-in-oil droplets can be triggered to form host condensates and capture proteins using UV irradiation^16^. In these experiments, we encapsulated a mixture of inactive nanostars (pre-annealed D_1,_ D_3_ and R_4_^apt^), strand D_2_^photo^, and SA by vortexing the sample with oil and surfactant mix using a benchtop vortexer^16,28^. Droplet samples were then transferred to Ibidi imaging chambers for observation as described in the Methods. As in the non-encapsulated experiments, we observed that the inactive nanostars (no UV irradiation) remain dispersed after encapsulation (example images are in Figure S7). Condensation begins only after UV irradiation, which cleaves the PC linker and activates the protected sticky end. As condensates fuse over time, a single condensate is visible in most droplets 3 hours after UV light irradiation (Figure S8); the condensates recruit SA client protein consistent with expectation, as shown by the very high correlation coefficients of the reporters for nanostars and proteins (Figure 3E). Figure 3F shows an example droplet and the condensate fluorescence intensity profile.

### Cotranscriptional control of nanostar condensation and protein recruitment

Another way to achieve kinetic control over condensation of our hybrid nanostars is to use the RNA output of synthetic genes to bind nanostars and enable their interactions (Figure 4A). First, we verified that the pre-annealed inactive nanostar core (D_1_, D_2_, and D_3_) can be activated by isothermal addition (at 37 °C) of the corresponding RNA strand in variants (C) and (D). Here spin-column purified RNA strands R_4_^b^ and R_4_^apt^ were added to separate samples including pre-annealed inactive nanostar cores, and we verified the formation of condensates; control samples including the inactive nanostar core alone do not yield condensates (Figure S9).

**Figure 4.**
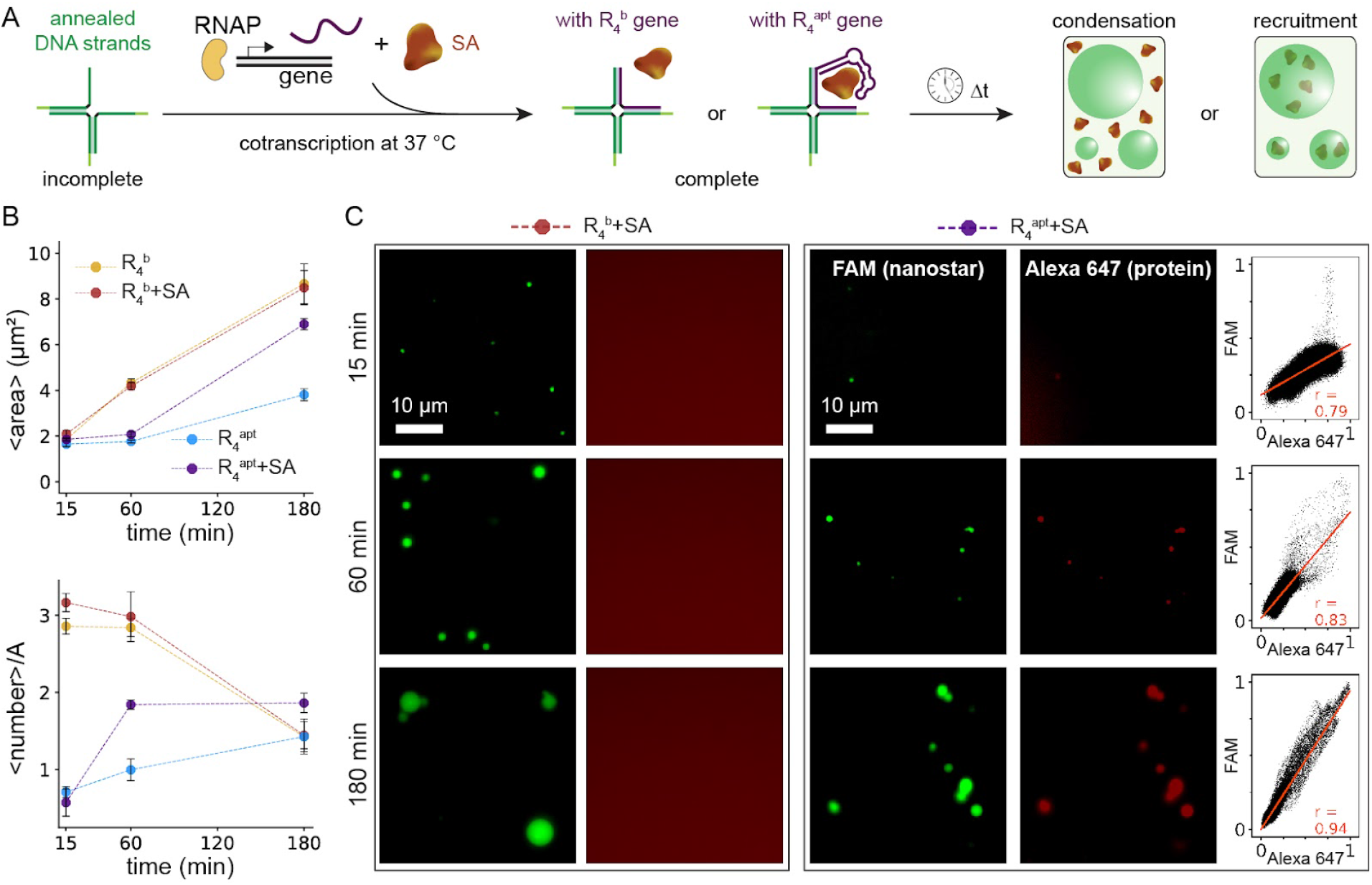
Cotranscriptional control of host condensate formation and protein capture. A) Schematic representation of cotranscriptional production of the RNA strands from synthetic genes at 37 °C and binding of the RNA strand that completes the inactive nanostar core. When the nanostar is complete (all the arms are double-stranded), they interact to form condensates; if the SA aptamer is present, SA protein is recruited to the dense phase. B) Plots of the average area and the number density of condensates up to 180 minutes in the absence (samples R_4_^b^ and R_4_^apt^) or presence of SA (samples R_4_^b^ + SA and R_4_^apt^ + SA). Dots correspond to the average of three experimental replicas (n = 3); error bars are the standard deviation of the mean. C) Fluorescence microscopy images of cotranscriptionally activated nanostars taken 15, 60, and 180 minutes after the start of transcription. SA recruitment does not occur for design (C) that includes R_4_^b^; SA is only recruited to condensates formed by design (D) that includes strand R_4_^apt^. The colocalization of FAM (condensates) and Alexa 647 (SA) reporters is confirmed by correlation plots (right).

Next, we tested whether inactive nanostars can be activated in a one pot transcription reaction at 37 °C from DNA templates (0.33 *μ*M) including the bacteriophage T7 promoter in the presence of T7 RNA polymerase (RNAP). We found that condensation occurs robustly both in the presence and in the absence of the client SA (Figure 4B and Figure S9). Figure 4C shows example fluorescence microscopy images of the resulting hybrid DNA-RNA condensates under transcription conditions, tracked up to 3 hours. The growth of condensate for design (D) is remarkably slower when compared to design (C). This indicates that the secondary structure of transcribed R_4_^apt^, as well as its length, may hinder isothermal binding to the inactive core. The slow increase in the number density for design (D) over 3 hours also indicates that fusion events may be rare when compared to the slow emergence of new condensates. Figure S10 includes box plots of condensate size in the aforementioned cases.

For design (C), which does not recruit SA, condensate growth and number density are comparable in the presence or absence of SA (Figure 4B). In contrast, the host condensates emerging from design (D) that includes R_4_^apt^ show faster growth and increase in number density when SA is present. The presence of SA may promote correct folding of R_4_^apt^ as it is transcribed, and thus facilitates nanostar activation. Colocalization of SA during transcription is confirmed by the correlation plots in Figure 4C. Finally, colocalization analysis of condensates formed in the absence and presence of SA is shown in Figure S11, reaffirming that only variant (D) is capable of protein recruitment.

### Control of protein recruitment with enzymes that produce and degrade RNA

Next, we sought to achieve transient protein recruitment to host condensates by combining catalytic production and degradation of RNA, which introduces a dissipative mechanism for condensate formation^30^. For this purpose, in addition to producing the RNA activating condensates cotranscriptionally using RNAP, we introduced degradation of the RNA hybridized to DNA using RNase H (Figure 5A). Having just demonstrated that the host nanostar design (D) can be activated cotranscriptionally (without RNA purification or annealing), we expected that RNase H-mediated RNA degradation would dissolve host condensates by degrading R_4_^apt^ hybridized to the DNA nanostar core DNA structure. Further, the RNase H level should regulate the timescale of dissolution. Because degradation of R_4_^apt^ also causes the release of SA from the nanostars, the RNase H level should also control the timescale of client protein localization.

**Figure 5.**
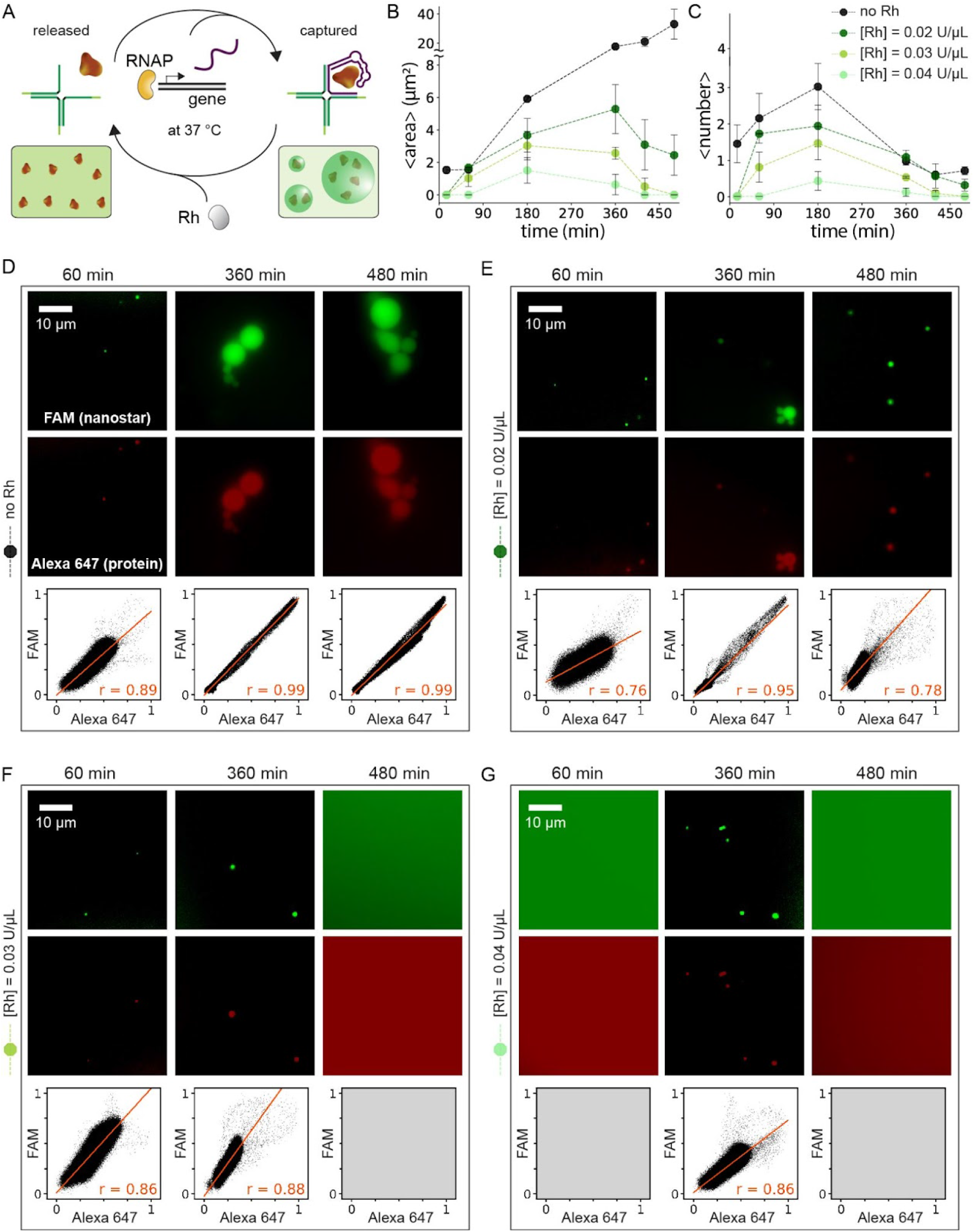
Transient capture and release of SA into host condensates mediated by enzymatic production and degradation of RNA. A) Schematic of the reactions occurring in a sample that includes inactive nanostar core, SA, gene, RNAP, and RNase H. B-C) The average condensate area and count increases when growth is triggered by the addition of the gene (black curve) in the absence of RNase H. By tuning the concentration of RNase H (0.00-0.04 U/μL), we can control the time required to dissolve condensates and thus protein recruitment (green curves). After an initial growth stage, RNase H-mediated degradation dominates resulting in the dissolution of RNA condensates and the release of SA. Dots correspond to the average of three experimental replicas (n = 3), and error bars represent the standard deviation of the mean. D-G) Fluorescence microscopy images of the sample for different RNase H levels; the size of condensates increases and SA is recruited only transiently at larger amounts of RNase H. The correlation of FAM and Alexa 647 signals indicates colocalization of condensates and SA during the experiment (bottom plots).

As a control experiment, we first tracked condensate growth and protein capture for up to 480 minutes in the absence of RNase H, and verified that the average condensate area continues to grow over time (Figure 5B, box plots are reported in Figure S12). The recruitment of SA is confirmed by the colocalization of reporters shown by correlation plots (left, Figure 5C). We then investigated how the temporal evolution of protein recruitment is affected at different levels of RNase H. Thus, in samples containing 2.5 μM inactive nanostar core, and gene (0.33 μM) producing R_4_^apt^, and a fixed level of RNAP (2.5% v/v), we varied the level of RNase H (0.02, 0.03 and 0.04 U/μL) while incubating the reaction at 37 °C (Figure 5B-C). In this range of RNase H levels, formation of condensates dominates for the first 3-4 hours of the reaction (no condensates formed, however, at 0.05 U/µL RNase H). The effects of RNA degradation become prominent after 4 hours: condensates dissolve, at a speed proportional to the amount of RNase H supplied, and result in delocalization of the SA client (Figure 5B-C and Figures S12-S13). We attribute the RNase H-dependent dominance of condensate dissolution to multiple phenomena that have been previously observed in DNA tile assembly^31^, including loss of RNAP activity over time and accumulation of transcription and degradation byproducts that form nanostar complexes lacking the correct structure for condensation.

Both condensate area and number density show a “pulsed” behavior, in which the duration and peak of the pulse inversely correlate with the amount of RNase H. Overall, these experiments show that autonomous, temporal recruitment of proteins to host condensates can be enabled by the interplay of RNA production and degradation. In our setup, transcription generates R_4_^apt^ that works as a fuel molecule supporting host condensate formation and protein recruitment. Degradation of R_4_^apt^ is a mechanism of dissipation of the fuel molecule, which drives the return to a uniform dispersed phase of the nanostars.

## Discussion

We have demonstrated methods to build nucleic acid host condensates for the recruitment of streptavidin (SA) as a model protein. The formation of condensates is driven by the interactions of DNA-RNA nanostructured motifs, which include an RNA aptamer that binds to SA. We illustrated two methods to trigger the formation of our host condensates and the localization of their target protein: we used UV light as a physical input, and transcription as an example biochemical reaction. Both approaches are amenable to fine tuning the growth rate of host condensates, for instance by dosing the UV irradiation time^16^, or by fine tuning the concentration of transcription components (template and RNAP)^32^. Further, we showed that the combination of reactions for RNA production and degradation leads to autonomous, transient formation of host condensates, and thus of protein recruitment.

The combination of DNA and RNA into a hybrid design has some advantages when compared to all-DNA or all-RNA nanostars. The DNA nanostar core proved to be a robust motif, because a range of variants generate condensates growing at comparable speeds. In our study we achieved UV-controllability of condensates by inserting a photolabile functional group in the DNA core, but different DNA modifications could render the condensates responsive to additional physical or chemical inputs^33^. Transcriptional control is contributed by the presence of an RNA molecule that is essential for condensation, in addition to having the role of recruiting a target ligand. RNA aptamers are an ever expanding class of molecules that can bind their target with high specificity and sensitivity^34^, and host condensates could potentially be used in the context of biosensing or toward the development of active biomaterials operating under physiological conditions. However, owing to the negatively-charged nature of nucleic acids, non-specific binding of nanostars to positively-charged protein may occur in the absence of aptamers, which points to a limitation of this approach. In addition to simply hosting and separating molecules, our condensates could also host catalytic reactions via RNA enzymes^34,35^.

Host condensates made with DNA or RNA motifs may be built through a variety of approaches, including those relying on long sequence repeats^36^, although DNA nanostars are particularly versatile. First, a variety of aptamers could be modularly appended to nanostar arms to localize specifically a diverse set of ligands, while excluding others depending on size and charge^37^. Another advantage of DNA nanostars is that their viscoelastic properties can be finely tuned by modifying the sticky end interaction energy and the number of arms^38^, in addition to being dependent on temperature and ionic conditions. This means that the local conditions of the dense phase could be optimized for specific guest molecules and reactions. Through sequence optimization, distinct coexisting DNA condensates can be spatially segregated, or made to interact to form compartments that include multiple subcompartments: this means DNA condensates could be used to host multi-step reactions and potentially enhance their yield^18,39,40^. Finally, the recruitment and release of guest molecules could be finely controlled through strand displacement reactions that could fold or disrupt the guest binding domains^41^.

## Methods

### Oligonucleotides and enzymes

All DNA strands were purchased from Integrated DNA Technologies (Coralville, IA). Oligonucleotides were standard desalted, except fluorescently labeled strands which were HPLC-purified. RNA strands were transcribed in-house and column-purified using the Monarch® RNA Cleanup Kit (New England Biolabs). The concentration of oligonucleotides was quantified by UV-vis spectrophotometry using a Thermo Fisher Scientific NanoDrop 2000 Spectrophotometer. All the oligonucleotides were resuspended in double-distilled H_2_O and kept at −20 °C for long-term storage. Sequences were either adapted from the literature or designed using Nupack^42^. Oligonucleotide sequences and modifications are provided in the SI Section 1. Streptavidin (Molecular Probes™ Streptavidin, Alexa Fluor™ 647 Conjugate) was purchased from Thermo Fisher Scientific. Transcription reactions were performed using the AmpliScribe T7-Flash RNA Polymerase (RNAP) enzyme. RNAse H (stock concentration: 5 U/μL) was ordered from Thermo Fisher Scientific. All the enzymes were stored at −20 °C.

### RNA production

The synthetic genes used to produce RNA were mixed in 1X transcription buffer (AmpliScribe 10X Reaction Buffer) and annealed from 90 °C to 25 °C at a rate of −1 °C/min. RNA was transcribed *in vitro* using the AmpliScribe T7-Flash Transcription Kit (Biosearch Technologies) for 3-4 hours, at 37 °C. The reaction was quenched by adding DNAse I (5% v/v) and incubating the sample at 37 °C for an additional 15-30 minutes. The Monarch® RNA Cleanup Kit (New England Biolabs) was used to purify transcribed RNA strands, whose concentrations were then measured by Nanodrop.

### Assembly of nanostars and condensate formation

Nanostars were prepared by mixing each strand at a final concentration of 2.5 μM in 12.5 mM MgCl_2_, 20 mM Tris-HCl, pH 8.0 buffer, using Eppendorf DNA Lobind® tubes. For experiments including SA, condensates were formed by annealing each strand at 2.7 μM, and SA was added immediately after annealing to reach at 2.5 μM final concentration of both nanostars and SA. Two variants of D_1_ were labeled either with FAM (D_1_^fam^, default in this study unless otherwise noted) or Cy3 (D_1_^cy3^). For visualizing condensates during fluorescence microscopy experiments, D_1_^fam^ was used at 0.5 μM, 10% or 20% molar ratio; D_1_^cy3^ was used at 2.5% molar ratio. To prepare the photoactivatable nanostars, the PC-modified strand (D_2_^photo^) was added isothermally at 25 °C after annealing. Annealing was performed by a Bio-Rad T100^TM^ Thermal Cycler, starting at 90 °C for all-DNA samples and at 72 °C for DNA-RNA samples and cooled down to room temperature at a rate of −1 °C/min. After annealing, samples were incubated at 25 °C unless otherwise noted. Throughout the study, the SA concentration was 2.5 μM (the same as the nanostar concentration).

For cotranscription experiments, the inactive nanostar core (D_1_+D_2_+D_3_) was annealed at an initial concentration of 5 μM in the aforementioned buffer, which then was added to the mixture of SA (if applicable) and cotranscription medium to reach a target concentration of 2.5 μM. The cotranscription mix contains 1X transcription buffer, 0.33 μM gene, 25 mM rNTPs, 10 mM MgCl_2_, and 5% (v/v) RNAP. For the SA capture and release experiments, the same cotranscription mix was used, to which we added the appropriate amount of RNAse H.

For the photoactivatable nanostars, strands D_1_, D_3_, as well the RNA strand (R_4_^b^ or R_4_^apt^), were first annealed from 72 °C to 25 °C, and the PC-modified strand (D_2_^photo^) was then added isothermally to the sample. Next, the sample was UV irradiated for 180 s at a roughly 4 cm distance by a 320 nm, 8 Watt light source (115 V, 60 Hz). Unless otherwise stated, all the experiments were done in triplicate (n = 3).

### Preparation of water-in-oil emulsion droplets

To generate emulsion droplets, we first prepared a 2% (w/w) mixture of surfactant (008 FluoroSurfactant, RAN Biotechnologies) and oil (FC-40 Fluorinert^TM^, Sigma-Aldrich). Using Eppendorf DNA Lobind® tubes, we combined 80 μL oil-surfactant mixture with 20 μL of pre-mixed SA and protected (inactive) hybrid nanostars including R_4_^apt^. Next, the sample was shaken for about 50 s using a bench vortexer to generate a large population of isolated, immiscible droplets of various sizes. The vortexed sample was equilibrated at room temperature for a few minutes prior to transferring an aliquot of the emulsion (milky fraction of the sample) into an Ibidi chamber (μ-Slide VI 0.4). The channel was sealed to prevent evaporation and to limit droplet motion due to pressure differences. The channel was irradiated with UV light for 180 s. All emulsion experiments were performed in triplicate (n = 3).

## Epifluorescence microscopy imaging

Prior to imaging, coverslips (Fisherbrand™, 60×22 mm^2^, 0.13 to 0.17 mm thick) were soaked in 5% (w/v) bovine serum albumin (BSA) for several minutes to prevent non-specific interactions of DNA on the glass surface. After BSA coating, the slides were then washed five times with distilled water and dried inside a fume hood. A small imaging chamber was prepared using a Parafilm (Fisher Scientific) square with a hole punched in the middle, taped to the washed coverslips by heating to 40-50 °C. After cooling the coverslip, 2 μL of sample was aliquoted inside the punched Parafilm area. The sample was covered with a second coverslip (Fisherbrand™, 22×22 mm^2^, 0.13-0.17 mm thick) to prevent evaporation of the sample during imaging. BSA coating prevents nanostar interactions with the glass surface, and limits surface-induced nucleation of condensates as well as their adhesion to the surface.

Samples were imaged with a Nikon Eclipse TI-E. Condensate images were acquired using a Nikon CFI Plan Apo Lambda 60X Oil objective, and Eclipse filter cubes with excitation wavelength of 448 nm for FAM-labeled nanostars and 646 nm for Alexa 647-labeled SA. Two separate channels, both with exposure times of 90 ms, were used to image the Alexa 647 and FAM dyes. For the majority of experiments, at least 10 multichannel images were acquired at each time point. Emulsion droplets were imaged using a Nikon Plan Fluor 20X/0.5 NA objective on a fixed field of view (time-lapse imaging).

### Image processing

We processed and analyzed all the microscopy images with custom Python scripts (available on Github: https://github.com/klockemel/Condensate-Detection). For each experimental condition, we randomly selected eight images out of a pool of at least 10 images and ran the script over their green channel, which ultimately calculated the condensate average area and number, along with its mean intensity. The normalized number of condensates reported in each figure corresponds to the average number of condensates measured over an area measuring 42.4×42.4 µm^2^ (corresponding to the size of images reported in the figures). This number was computed from the total number of condensates measured over eight fields of view (144.8×144.8 µm^2^), and averaged over three experimental replicates. Colocalization analysis was performed using a Fiji ImageJ plugin, Just Another Colocalization Plugin (JACoP)^43,44^, available at https://imagej.net/plugins/jacop. The pixel intensities of both channels (*X*_*i*_) were obtained, normalized as *x*_*i*_ = (*X*_*i*_ − *X*_*min*_)/(*X*_*max*_ − *X*_*min*_), and plotted using Python. Pearson’s correlation coefficient corresponding to each microscopy snapshot shown in the figures was calculated using Python’s Scipy library.

### Native PAGE experiments

Native polyacrylamide gels (15%) were prepared with TBE (10X), 30% ammonium persulfate (APS, 0.18% v/v), and tetramethylethylenediamine (TEMED, 0.11% v/v). Gels were cast in 10×10 cm^2^, 1.5 mm thick disposable minigel cassettes and allowed to polymerize for at least 30 minutes before electrophoresis. The gel was incubated with a running buffer (1X TBE solution, pH 8.0) for 30 minutes at 25 °C. Sample volumes of 10 μL were combined with 1 μL of 6X Orange DNA Loading Dye and then loaded directly into the gel. Native PAGE was run in an electrophoresis unit at 25 °C using 1X TBE buffer at pH 8.0 and a constant voltage of 120 V for 150 minutes. Gels were stained with SYBR Gold for 20 minutes and scanned using a ChemiDoc^TM^ MP imaging system (Bio-Rad).

## Supporting information

SupportingInformation

## Acknowledgements

We thank Dino Osmanovic, Olivia Zou, and Deborah Fygenson for helpful discussions and advice. This research was supported by the National Science Foundation (NSF) grant FMRG: Bio award 2134772, and by the Sloan Foundation through award G-2021-16831 to EF. DS is supported by a postdoctoral fellowship from the Associazione Italiana per la Ricerca sul Cancro (AIRC).

## References

1. Elani Y, Law RV, Ces O. Vesicle-based artificial cells as chemical microreactors with spatially segregated reaction pathways. Nat Commun. 2014;5:5305.

2. Hwang ET, Lee S. Multienzymatic Cascade Reactions via Enzyme Complex by Immobilization. ACS Catal. 2019;9(5):4402–4425.

3. Gomes E, Shorter J. The molecular language of membraneless organelles. J Biol Chem. 2019;294(18):7115–7127.

4. Banani SF, Lee HO, Hyman AA, Rosen MK. Biomolecular condensates: organizers of cellular biochemistry. Nat Rev Mol Cell Biol. 2017;18(5):285–298.

5. Lyon AS, Peeples WB, Rosen MK. A framework for understanding the functions of biomolecular condensates across scales. Nat Rev Mol Cell Biol. 2021;22(3):215–235.

6. Reinkemeier CD, Girona GE, Lemke EA. Designer membraneless organelles enable codon reassignment of selected mRNAs in eukaryotes. Science. 2019;363(6434). 10.1126/science.aaw2644

7. Kojima T, Takayama S. Membraneless Compartmentalization Facilitates Enzymatic Cascade Reactions and Reduces Substrate Inhibition. ACS Appl Mater Interfaces. 2018;10(38):32782–32791.

8. Lim S, Clark DS. Phase-separated biomolecular condensates for biocatalysis. Trends Biotechnol. 2024;42(4):496–509.

9. Schuster BS, Reed EH, Parthasarathy R, et al. Controllable protein phase separation and modular recruitment to form responsive membraneless organelles. Nat Commun. 2018;9(1):2985.

10. Abraham GR, Chaderjian AS, N Nguyen AB, Wilken S, Saleh OA. Nucleic acid liquids. Rep Prog Phys. 2024;87(6). 10.1088/1361-6633/ad4662

11. Udono H, Gong J, Sato Y, Takinoue M. DNA Droplets: Intelligent, Dynamic Fluid. Adv Biol (Weinh). 2023;7(3):e2200180.

12. Biffi S, Cerbino R, Bomboi F, et al. Phase behavior and critical activated dynamics of limited-valence DNA nanostars. Proc Natl Acad Sci U S A. 2013;110(39):15633–15637.

13. Jeon B-J, Nguyen DT, Abraham GR, Conrad N, Fygenson DK, Saleh OA. Salt-dependent properties of a coacervate-like, self-assembled DNA liquid. Soft Matter. 2018;14(34):7009–7015.

14. Agarwal S, Osmanovic D, Dizani M, Klocke MA, Franco E. Dynamic control of DNA condensation. Nat Commun. 2024;15(1):1915.

15. Do S, Lee C, Lee T, Kim D-N, Shin Y. Engineering DNA-based synthetic condensates with programmable material properties, compositions, and functionalities. Sci Adv. 2022;8(41):eabj1771.

16. Agarwal S, Dizani M, Osmanovic D, Franco E. Light-controlled growth of DNA organelles in synthetic cells. Interface Focus. 2023;13(5):20230017.

17. Zhao Q-H, Cao F-H, Luo Z-H, Huck WTS, Deng N-N. Photoswitchable molecular communication between programmable DNA-based artificial membraneless organelles. Angew Chem Weinheim Bergstr Ger. 2022;134(14). 10.1002/ange.202117500

18. Leathers A, Walczak M, Brady RA, et al. Reaction-Diffusion Patterning of DNA-Based Artificial Cells. J Am Chem Soc. 2022;144(38):17468–17476.

19. Stewart JM, Li S, Tang A, et al. Modular RNA motifs for orthogonal phase separated compartments. BioRxiv. 2023.

20. Fabrini G, Nuccio SP, Stewart JM, et al. Co-transcriptional production of programmable RNA condensates and synthetic organelles. BioRxiv. 2023.

21. Jain A, Cheng K. The principles and applications of avidin-based nanoparticles in drug delivery and diagnosis. J Control Release. 2017;245:27–40.

22. Silverman AD, Karim AS, Jewett MC. Cell-free gene expression: an expanded repertoire of applications. Nat Rev Genet. 2020;21(3):151–170.

23. Leppek K, Stoecklin G. An optimized streptavidin-binding RNA aptamer for purification of ribonucleoprotein complexes identifies novel ARE-binding proteins. Nucleic Acids Res. 2014;42(2):e13.

24. Nguyen DT, Saleh OA. Tuning phase and aging of DNA hydrogels through molecular design. Soft Matter. 2017;13(32):5421–5427.

25. Bai X, Li Z, Jockusch S, Turro NJ, Ju J. Photocleavage of a 2-nitrobenzyl linker bridging a fluorophore to the 5′ end of DNA. Proceedings of the National Academy of Sciences. 2003;100(2):409–413.

26. Weitz M, Kim J, Kapsner K, Winfree E, Franco E, Simmel FC. Diversity in the dynamical behaviour of a compartmentalized programmable biochemical oscillator. Nat Chem. 2014;6(4):295–302.

27. Dupin A, Simmel FC. Signalling and differentiation in emulsion-based multi-compartmentalized in vitro gene circuits. Nat Chem. 2019;11(1):32–39.

28. Agarwal S, Klocke MA, Pungchai PE, Franco E. Dynamic self-assembly of compartmentalized DNA nanotubes. Nat Commun. 2021;12(1):1–13.

29. Conrad N, Chang G, Fygenson DK, Saleh OA. Emulsion imaging of a DNA nanostar condensate phase diagram reveals valence and electrostatic effects. J Chem Phys. 2022;157(23):234203.

30. Del Grosso E, Franco E, Prins LJ, Ricci F. Dissipative DNA nanotechnology. Nat Chem. 2022;14(6):600–613.

31. Agarwal S, Franco E. Enzyme-driven assembly and disassembly of hybrid DNA–RNA nanotubes. J Am Chem Soc. 2019.

32. Sorrentino D, Ranallo S, Nakamura E, Franco E, Ricci F. Synthetic Genes For Dynamic Regulation Of DNA-Based Receptors. Angew Chem Int Ed Engl. 2024;63(17):e202319382.

33. Madsen M, Gothelf KV. Chemistries for DNA Nanotechnology. Chem Rev. 2019;119(10):6384–6458.

34. Famulok M, Hartig JS, Mayer G. Functional aptamers and aptazymes in biotechnology, diagnostics, and therapy. Chem Rev. 2007;107(9):3715–3743.

35. Zhang J, Lau MW, Ferré-D’Amaré AR. Ribozymes and riboswitches: modulation of RNA function by small molecules. Biochemistry. 2010;49(43):9123–9131.

36. Liu W, Deng J, Song S, Sethi S, Walther A. A facile DNA coacervate platform for engineering wetting, engulfment, fusion and transient behavior. Commun Chem. 2024;7(1):100.

37. Nguyen DT, Jeon B-J, Abraham GR, Saleh OA. Length-Dependence and Spatial Structure of DNA Partitioning into a DNA Liquid. Langmuir. 2019;35(46):14849–14854.

38. Conrad N, Saleh OA, Fygenson DK. Towards rational design of power-law rheology via DNA nanostar networks. arXiv [cond-mat.soft]. 2023.

39. Malouf L, Tanase DA, Fabrini G, et al. Sculpting DNA-based synthetic cells through phase separation and phase-targeted activity. Chem. 2023;2023.03.17.533162. 10.1101/2023.03.17.533162

40. Jeon B-J, Nguyen DT, Saleh OA. Sequence-Controlled Adhesion and Microemulsification in a Two-Phase System of DNA Liquid Droplets. J Phys Chem B. 2020;124(40):8888–8895.

41. Lloyd J, Tran CH, Wadhwani K, Cuba Samaniego C, Subramanian HKK, Franco E. Dynamic Control of Aptamer-Ligand Activity Using Strand Displacement Reactions. ACS Synth Biol. 2018;7(1):30–37.

42. Zadeh JN, Steenberg CD, Bois JS, et al. NUPACK: Analysis and design of nucleic acid systems. J Comput Chem. 2011;32(1):170–173.

43. Bolte S, Cordelières FP. A guided tour into subcellular colocalization analysis in light microscopy. J Microsc. 2006;224(Pt 3):213–232.

44. Costes SV, Daelemans D, Cho EH, Dobbin Z, Pavlakis G, Lockett S. Automatic and quantitative measurement of protein-protein colocalization in live cells. Biophys J. 2004;86(6):3993–4003.

