## SupportingInformation for "Protein recruitment to dynamic DNA-RNA host condensates"

### 1. Nucleic acid sequences

The DNA nanostars were modified based on the previous works[1–3] and tested using NUPACK[4,5] along with the aptamer domain. The T7 RNA Polymerase (RNAP) promoter is included in the template and non-template strands (blue and red domains) to facilitate transcription and minimize unwanted byproducts in the reaction[6]. The spacers (i.e., 'TT' and 'UU' for DNA and RNA strands, correspondingly) in the junction and sticky ends (i.e., 'GCGC') are bold.

***Table S1.*** *Oligonucleotide sequences of nanostars and gene templates. The ordinary DNA strands ($D_i$), along with the dyed ones ($D_1^{fam}$ and $D_1^{cy3}$), were taken from Agarwal et al.[2], and $R_4^b$ was adapted from the same reference by eliminating the sticky end and converting the strand to RNA. $R_4^{apt}$ is the elongation of $R_4^b$ which contains an additional domain corresponding to the streptavidin (SA) aptamer. The PC-modified strand was utilized from our previous work[1].*

| Name | 5' – sequence – 3' |
|---|---|
| $D_1$ | **GCGC**CAGTGAGGACGGAAG**TT**TGTCGTAGCATCGCACC |
| $D_1^{fam}$<br>$D_1^{cy3}$ | **fam**-CAGTGAGGACGGAAG**TT**TGTCGTAGCATCGCACC<br>**cy3**-CAGTGAGGACGGAAG**TT**TGTCGTAGCATCGCACC |
| $D_2$ | **GCGC**CAACCACGCCTGTCCA**TT**ACTTCCGTCCTCACTG |
| $D_2^{photo}$ | **CGCACCAAAGGT**/iSpPC/**GCGC**CAACCACGCCTGTCCA**TT**ACTTCCGTCCTCACTG |
| $D_3$ | **GCGC**CCATGGTCCCAAGTGA**TT**TGGACAGGCGTGGTTG |
| $D_4$ | **GCGC**GGTGCGATGCTACGAC**TT**TCACTTGGGACCATGG |
| $D_4^b$ | GGTGCGATGCTACGAC**TT**TCACTTGGGACCATGG |
| $R_4^b$ | GGUGCGAUGCUACGAC**UU**UCACUUGGGACCAUGG |
| NT-$R_4^b$ | **GTACGTAATACGACTCACTATA**GGTGCGATGCTACGAC**TT**TCACTTGGGACCATGG |
| T-$R_4^b$ | CCATGGTCCCAAGTGA**AA**GTCGTAGCATCGCACC**TATAGTGAGTCGTATTACGTAC** |
| $R_4^{apt}$ | **G**AUGCGGCCGCCGACCAGAAUCAUGCAAGUGCGUAAGAUAGUCGCGGGUCGGCGGCC<br>GCAU**C**GGUGCGAUGCUACGAC**UU**UCACUUGGGACCAUGG |
| NT-$R_4^{apt}$ | **GTACGTAATACGACTCACTATAG**ATGCGGCCGCCGACCAGAATCATGCAAGTGCGTA<br>AGATAGTCGCGGGTCGGCGGCCGCAT**C**GGTGCGATGCTACGAC**TT**TCACTTGGGACC<br>ATGG |
| T-$R_4^{apt}$ | CCATGGTCCCAAGTGA**AA**GTCGTAGCATCGCACC**G**ATGCGGCCGCCGACCCGCGACT<br>ATCTTACGCACTTGCATGATTCTGGTCGGCGGCCGCAT**CTATAGTGAGTCGTATTAC**<br>**GTAC** |

### 2. Supplementary figures

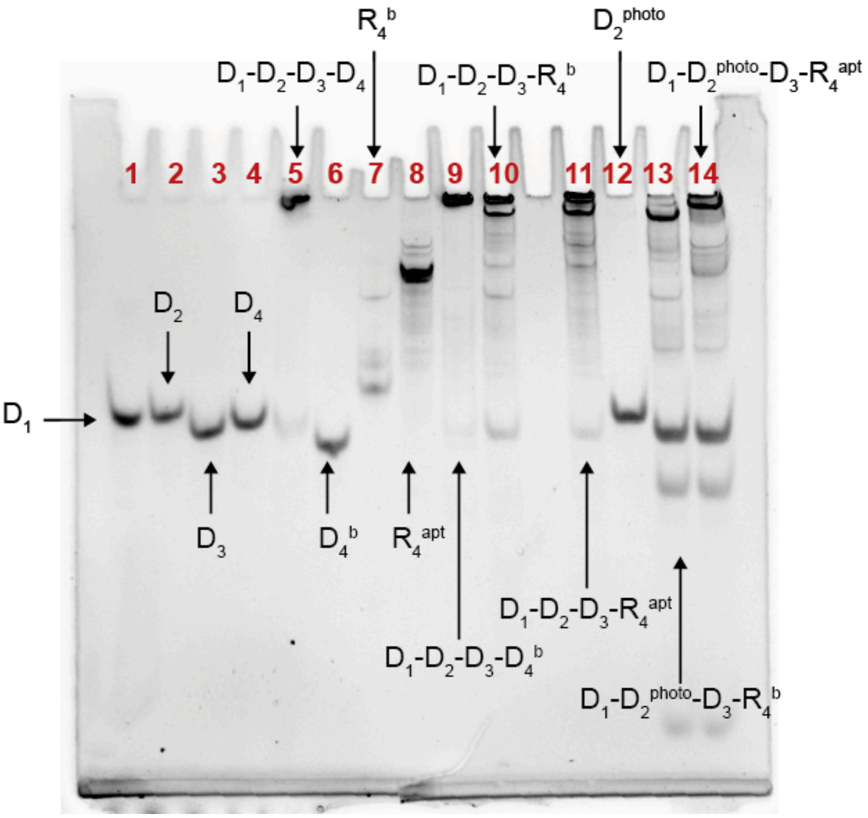


***Figure S1.*** *Native PAGE gel (15% polyacrylamide) showing* strand interactions and nanostar folding. *Each solution was prepared in a 20 mM Tris-HCl, 12.5 mM $MgCl_2$ buffer and incubated for 6 hours prior to running the gel. The gel was run at 25°C, 110 V, for 150 minutes in 1X TBE buffer, pH 8.0, stained for 20 min with 1X SYBR-Gold, and then imaged using the ChemiDoc MP imaging system (Bio-Rad).* ***Lane 1-4:*** *DNA strands for design (A).* ***Lane 5:*** *Annealed sample with design (A).* ***Lane 6:*** *Blunt-ended DNA strand ($D_4^b$).* ***Lane 7:*** *Blunt-ended RNA strand ($R_4^b$).* ***Lane 8:*** *RNA strand modified to contain the aptamer site for protein binding ($R_4^{apt}$).* ***Lanes 9, 10, 11:*** *Annealed samples of design (B), design (C), and design (D), respectively. A significant fraction of the complexes, corresponding to condensates, remain in the wells.* ***Lane 12:*** *PC-modified strand ($D_2^{photo}$).* ***Lanes 13-14:*** *Annealed samples of incomplete designs (C) and design (D), to which $D_2^{photo}$ is added isothermally; the sample is irradiated for 180 s.*

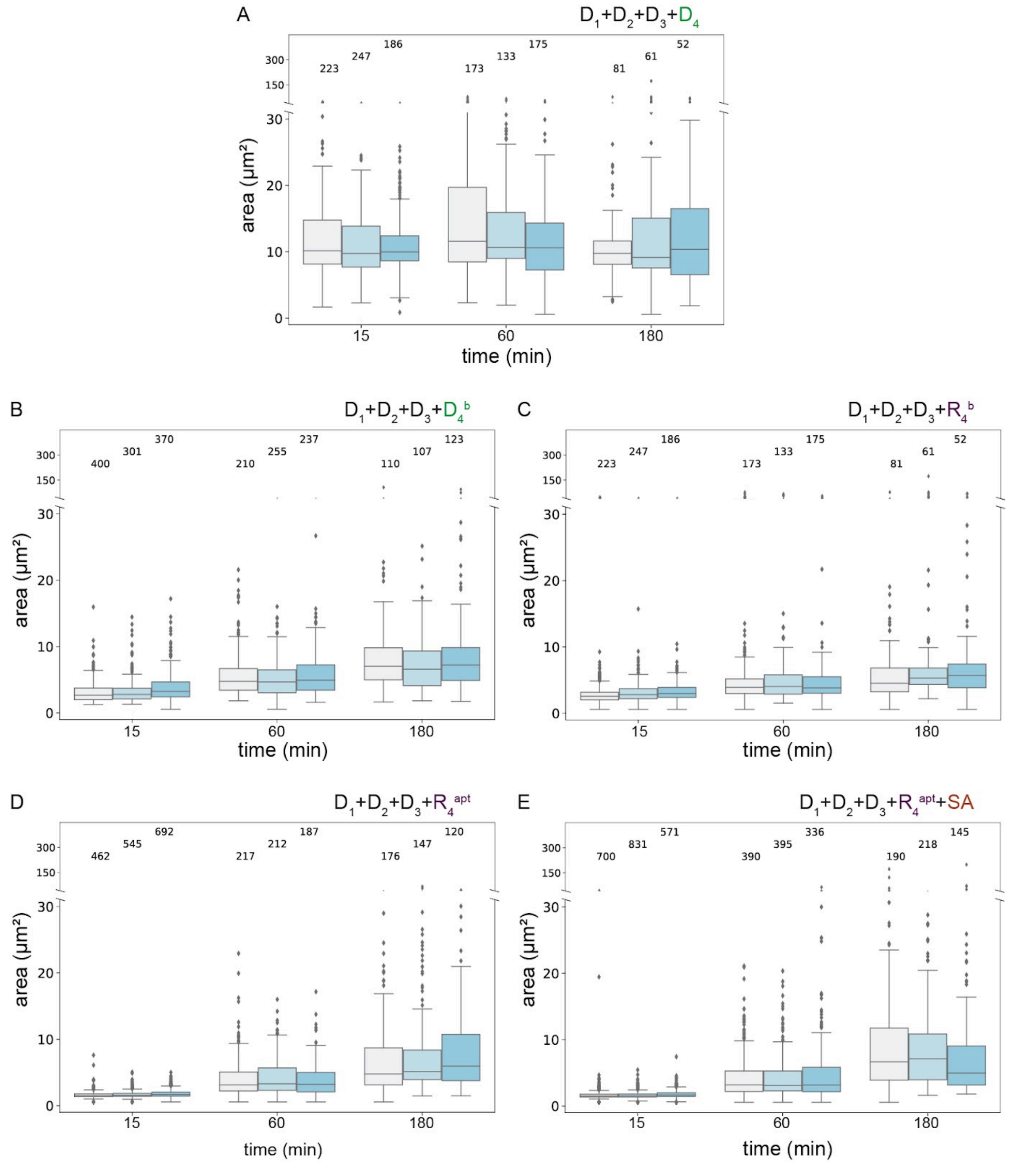


***Figure S2. Condensation of DNA-RNA 4-arm nanostars: box plots of the condensate size distribution for experiments reported in Figure 2 of the manuscript.*** *Images were taken 15, 60, and 180 minutes after the end of annealing. Box plots for the: A) experiments using design (A) that includes* $D_1$*,* $D_2$*,* $D_3$*, and* $D_4$*; B) experiments including design (B) that includes* $D_1$*,* $D_2$*,* $D_3$*, and* $D_4^b$*; C) experiments including design (C) that uses* $D_1$*,* $D_2$*,* $D_3$*, and* $R_4^b$*; D) experiments including design (D) that is composed of* $D_1$*,* $D_2$*,* $D_3$*, and* $R_4^{apt}$*; E) experiments including design (D) in the presence of SA. The distributions pool the data of triplicate, and the number of captured condensates associated with each time point is annotated on top of the plots.*

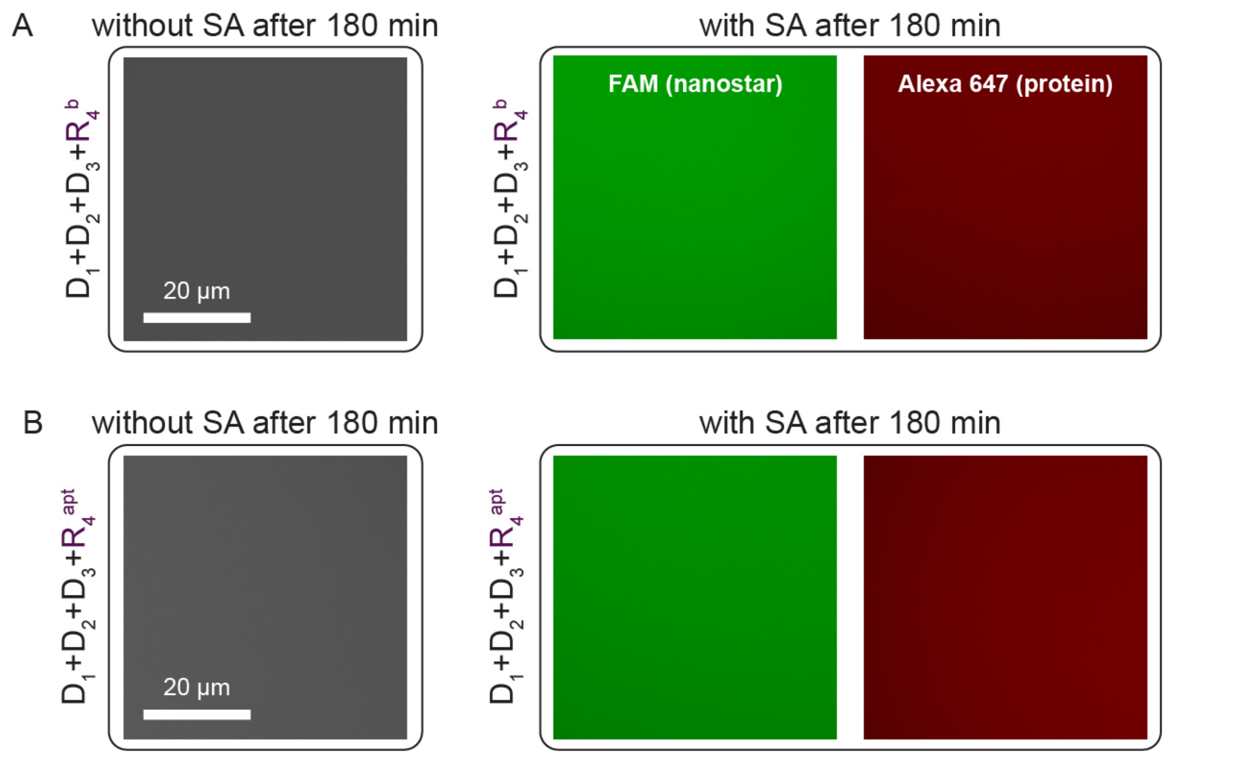


***Figure S3. UV irradiation is essential to trigger condensation of nanostars with PC-modified strand.*** *Representative fluorescence microscopy images of inactive DNA-RNA nanostars, using design (C) and design (D), in the absence of UV-irradiation after 180 minutes. No condensates are formed with A) $R_4^b$, and B) $R_4^{apt}$ neither in the absence (left) nor in the presence (right) of SA. Here, when SA was absent in the experiment, cy3 was used to label nanostars.*

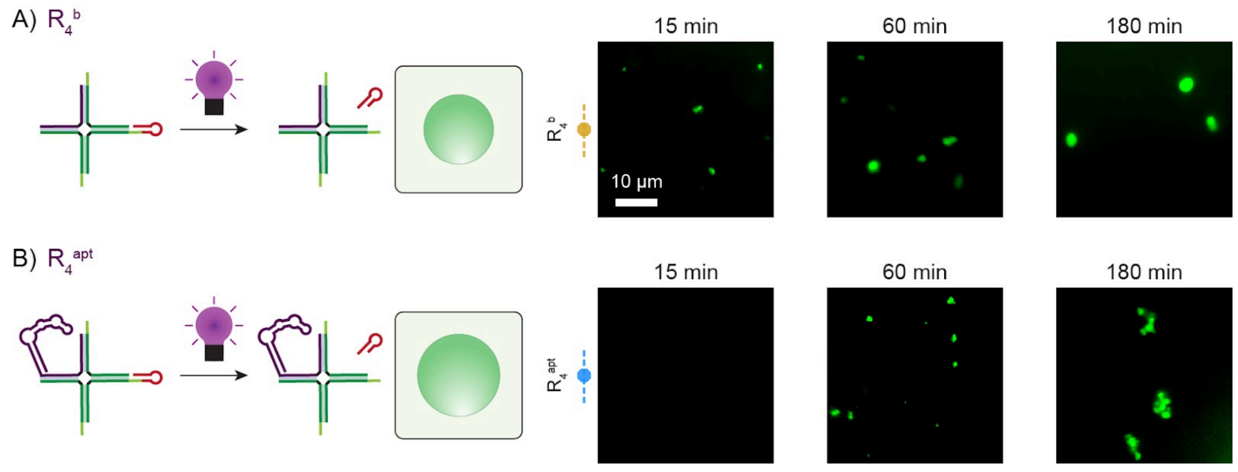


***Figure S4. Photoactivatable hybrid DNA-RNA nanostar motifs.*** *A) Left: Schematic of a 4-arm DNA-RNA nanostar, variant (C), activated by UV light ($t_{UV}$ = 180 s) in the presence of the PC-modified strand in the nanostars design. The UV irradiation activates the sticky ends and allows the nanostars to condense. Here, the RNA strand ($R_4^b$) is blunt-ended (without sticky end). Right: representative fluorescence microscopy images over time for UV-activated nanostars phase separating into condensates. B) Left: 4-arm DNA-RNA nanostar, variant (D), modified to contain a PC linker at the end of one DNA strand. The RNA strand here ($R_4^{apt}$) includes an aptamer domain for protein binding. Right: representative fluorescence microscopy images of activated nanostars condensing over time. All the images shown were taken at 15, 60, and 180 minutes after activation.*

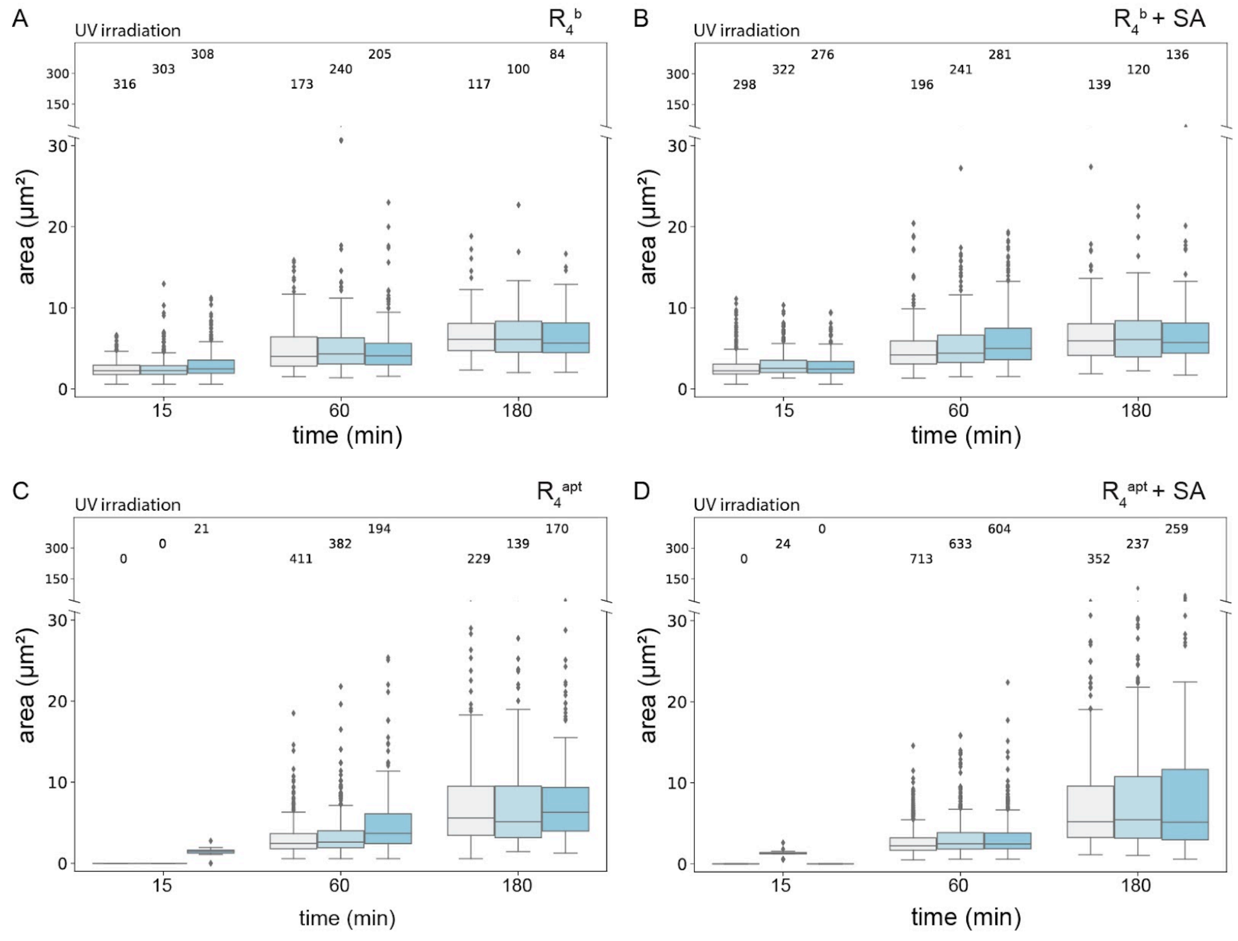


***Figure S5. Condensation of DNA-RNA 4-arm nanostars activated via UV irradiation: box plots of the condensate size distribution for experiments reported in Figure 3 of the manuscript.*** *Images were taken 15, 60, 180 minutes after UV irradiation for 180 s. Box plots for the: A) experiments using design (C) that includes* $D_1$*,* $D_2$*,* $D_3$*, and* $R_4^b$ *in the absence of SA; B) experiments using design (C) that includes* $D_1$*,* $D_2$*,* $D_3$*,* $R_4^b$ *in the presence of SA; C) experiments including design (D) that uses* $D_1$*,* $D_2$*,* $D_3$*, and* $R_4^{apt}$ *in the absence of SA; D) experiments including variant (D) that uses* $D_1$*,* $D_2$*,* $D_3$*,* $R_4^{apt}$ *in the presence of SA. The distributions pool the data of triplicate, and the number of captured condensates associated with each time point is given on top of the plots.*

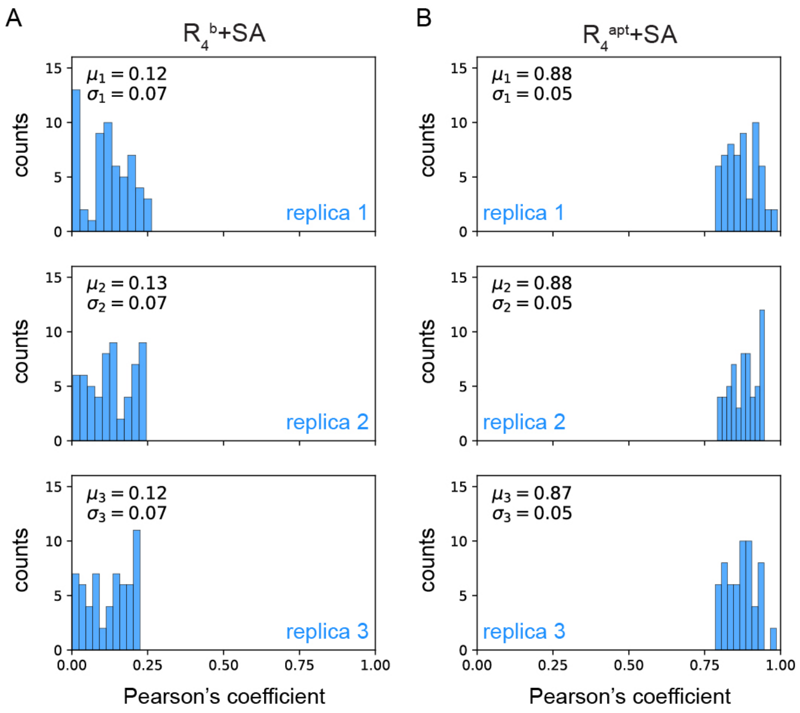


***Figure S6. Colocalization analysis of protein recruitment via UV irradiation.*** *Statistical distribution of Pearson's coefficient (r) of condensates, in triplicate, 180 minutes after UV activation ($t_{UV}$ = 180 s) in the cases of: A) $R_4^b$ in the presence of SA; B) $R_4{}^{apt}$ in the presence of SA. Values close to 0 for r (A) indicate low colocalization of the FAM and Alexa 647 fluorophores (nanostars and SA, respectively), suggesting no protein recruitment; while r values close to 1 (B) indicate high colocalization, thus demonstrating protein recruitment. The mean value (μ) and standard deviation (σ) of r for each replica are calculated from a random selection of 60 condensates.*

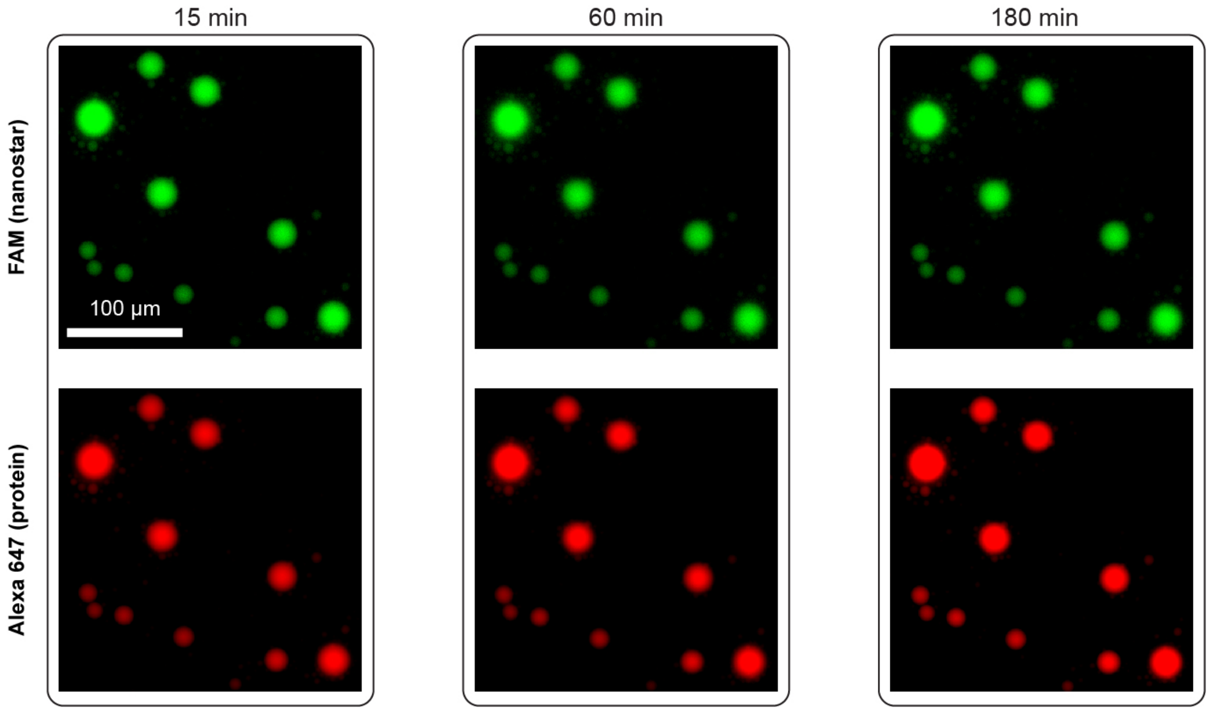


***Figure S7. Condensates cannot form inside water-in-oil droplets in the absence of UV irradiation.*** *Representative microscopy images of the encapsulated samples, containing inactive hybrid nanostars with aptamer and SA, at 15, 60, and 180 minutes after encapsulation. No UV light was irradiated to the samples, and the experiment was repeated three times (n = 3).*

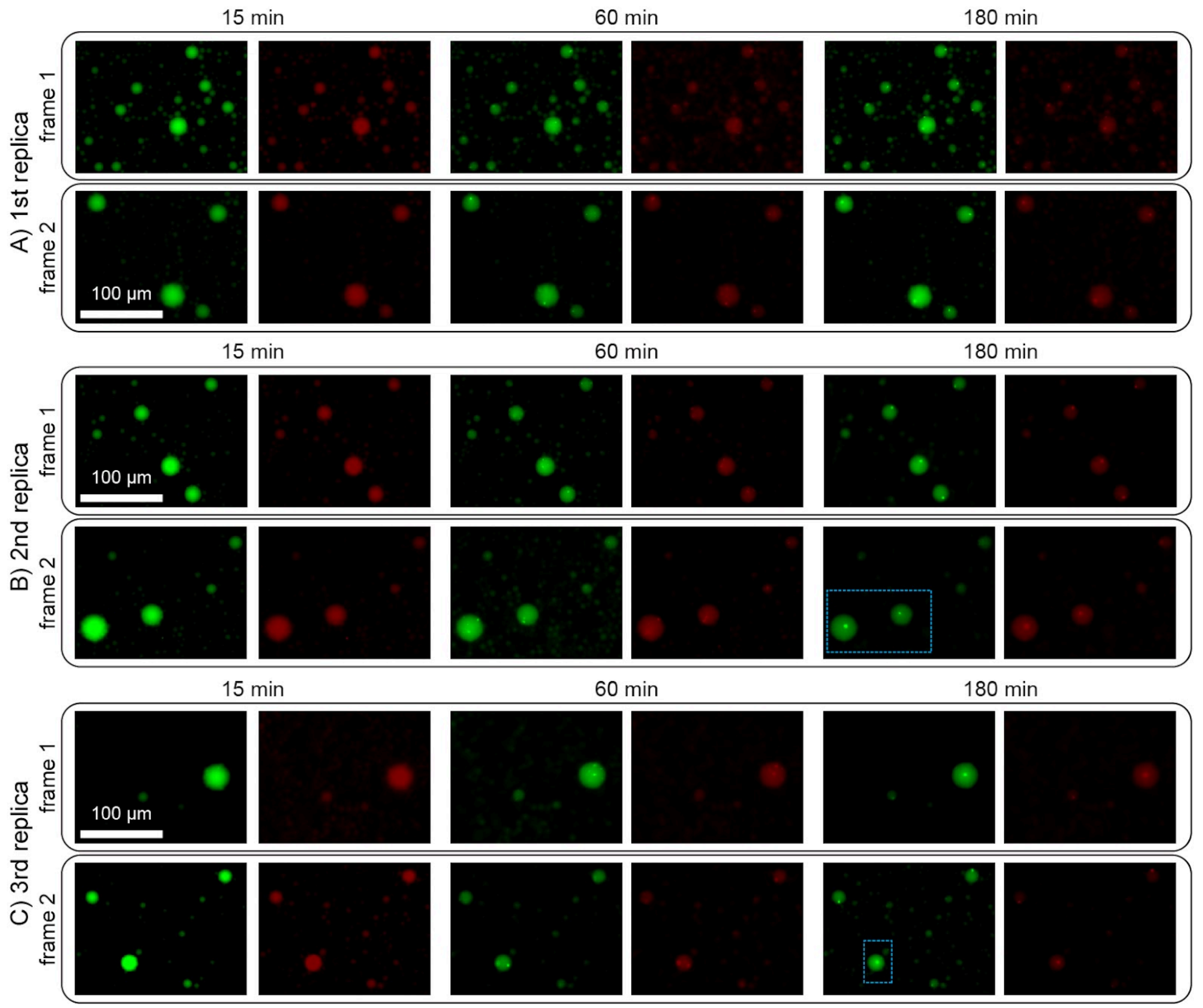


***Figure S8. Fluorescent microscopy images of light-controlled activation of condensates which capture protein inside water-in-oil droplets.*** *Microscopy images of the condensation of hybrid nanostars that have aptamer binding to SA at 15, 60, and 180 minutes after UV irradiation ($t_{UV}$ = 180 s). Each panel represents one replica. The dashed boxes in (B) and (C) represent the frames shown in (E) and (F) in Figure 3, respectively. At each time point, the left and right snapshots are FAM (nanostar) and Alexa 647 (protein), correspondingly.*

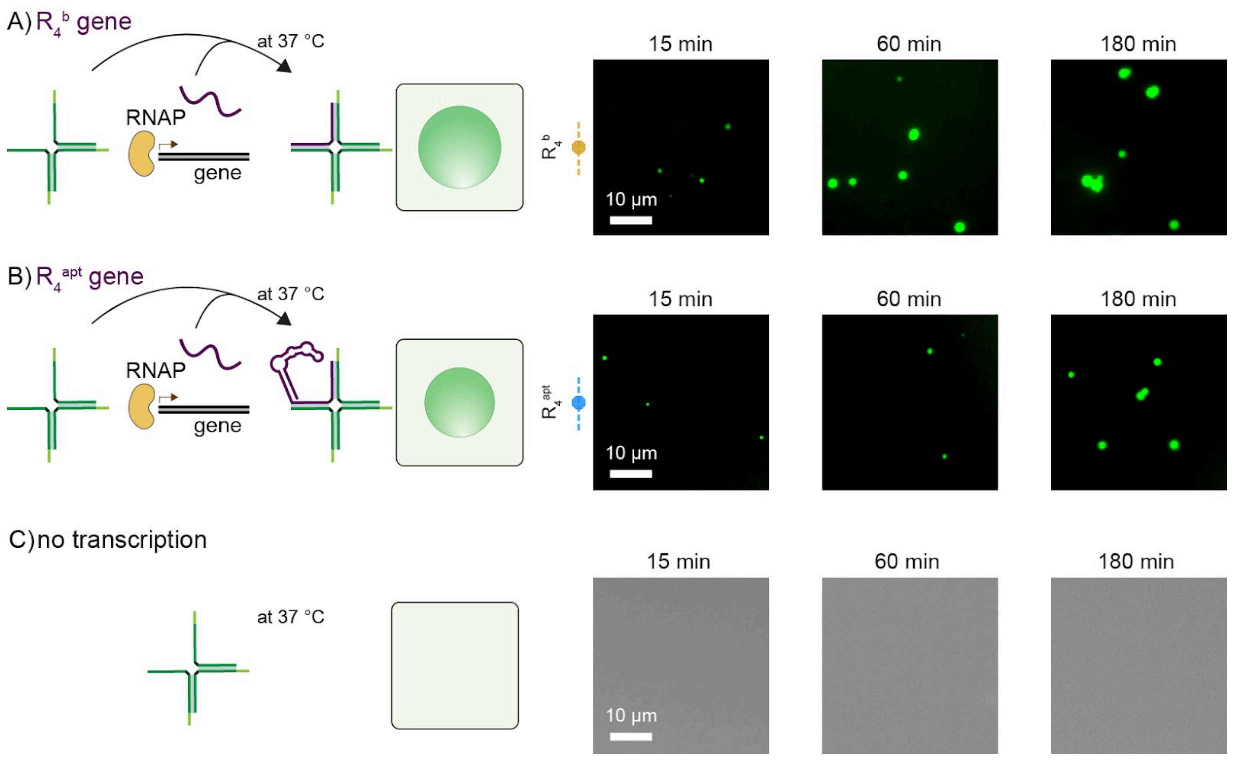


***Figure S9. Cotranscriptional activation of hybrid DNA-RNA nanostar motifs.*** *A) Left: Cotranscriptional activation of incomplete nanostars using a synthetic gene that transcribes a blunt-ended RNA strand. Right: Representative fluorescence microscopy images showing condensate growth over time under the cotranscription condition. B) Cotranscriptional activation of the hybrid nanostars using a synthetic gene (blue) that transcribes an aptamer-modified RNA strand for protein recruitment (purple). C) the core structure without RNA cotranscription does not form condensate at 37 °C (n = 1). Right: Representative fluorescence microscopy images, labeled with cy3, showing no condensation in the absence of cotranscription reaction.*

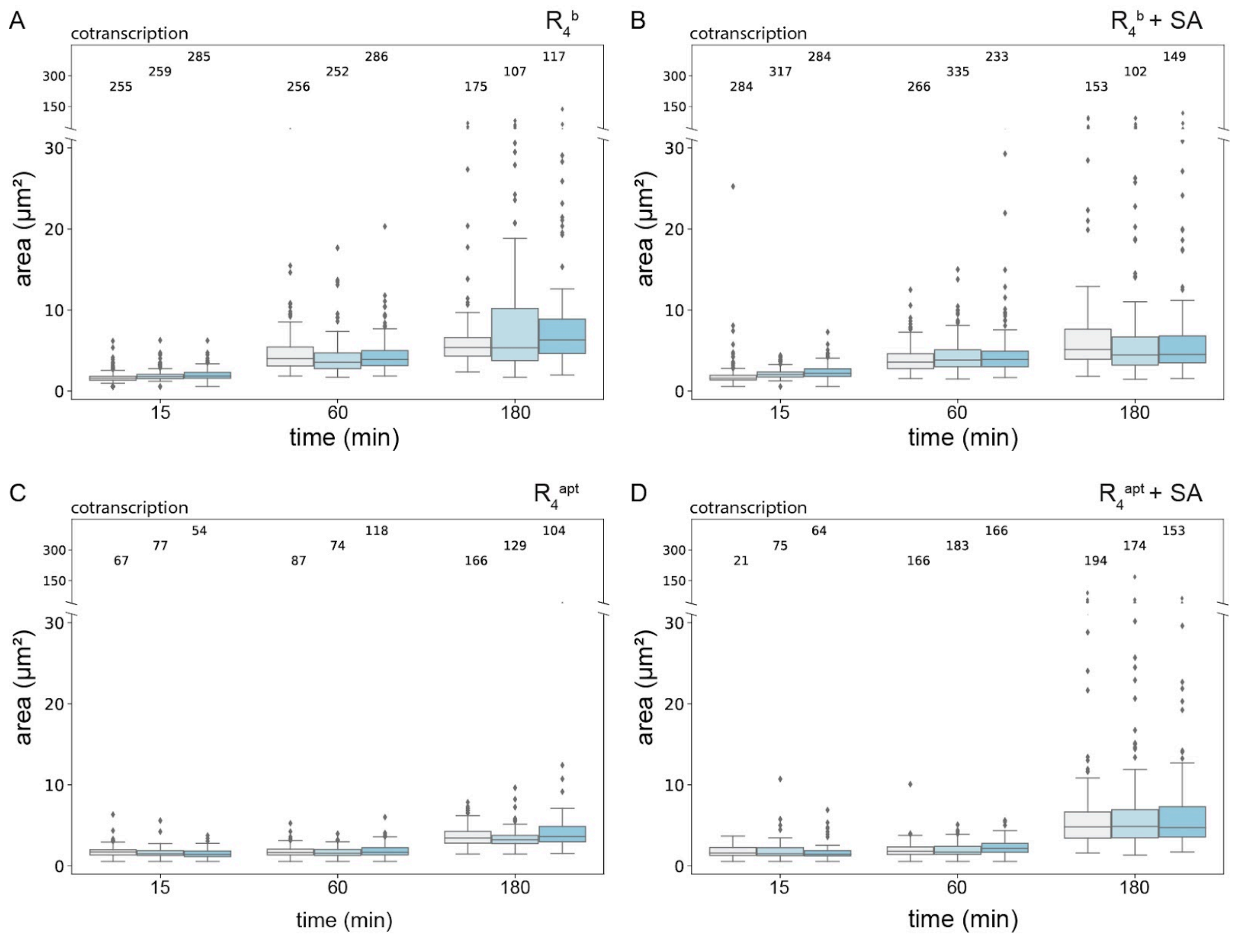


***Figure S10. Condensation of DNA-RNA 4-arm nanostars activated via cotranscriptional production of the RNA strand: box plots of the condensate size distribution for experiments reported in Figure 4 of the manuscript.*** *Images were taken 15, 60, and 180 minutes after transcription started. Box plots for the: A) experiments using design (C) that includes* $D_1$, $D_2$, $D_3$, *and* $R_4^b$ *in the absence of SA; B) experiments using design (C) that includes* $D_1$, $D_2$, $D_3$, *and* $R_4^b$ *in the presence of SA; C) experiments including design (D) that uses* $D_1$, $D_2$, $D_3$, $R_4^{apt}$ *in the absence of SA; D) experiments including design (D) that uses* $D_1$, $D_2$, $D_3$, $R_4^{apt}$ *in the presence of SA. The distributions pool the data of triplicate, and the number of captured condensates associated with each time point is written on top of the plots.*

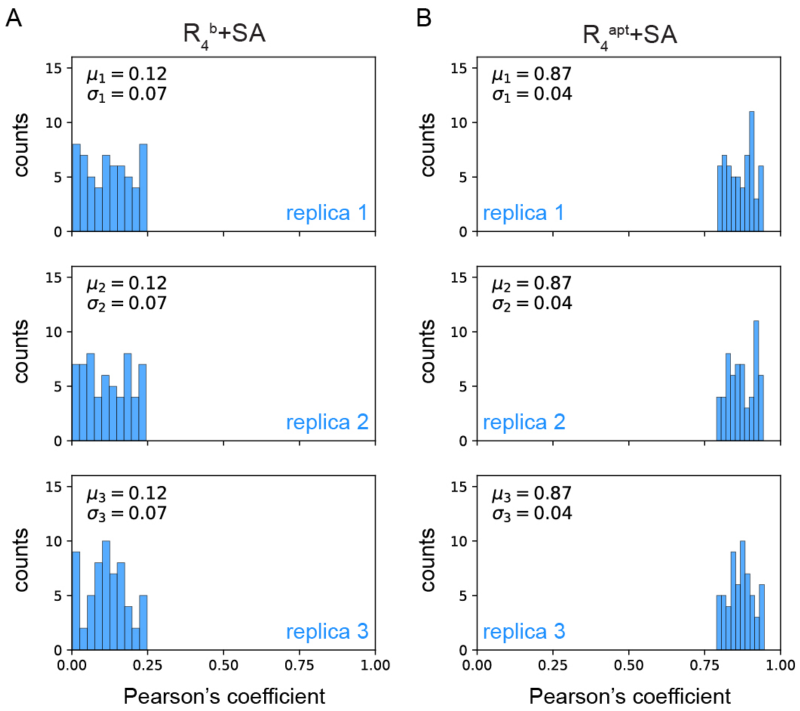


***Figure S11. Colocalization analysis of protein recruitment via cotranscriptional production of the RNA strand.*** *Statistical distribution of Pearson's coefficient (r), for three replicas, of condensates 180 minutes after the start of cotranscription in the cases of: A) $R_4^b$ in the presence of SA; B) $R_4^{apt}$ in the presence of SA. The r values close to 0 (A) indicate low colocalization of the FAM (nanostar) and Alexa 647 (protein) fluorophores and no protein recruitment; whereas the r values close to 1 (B) indicate high colocalization, corroborating protein recruitment. The mean values of r (μ) and its standard deviations (σ) for each replica are obtained from analyzing a random selection of 60 condensates.*

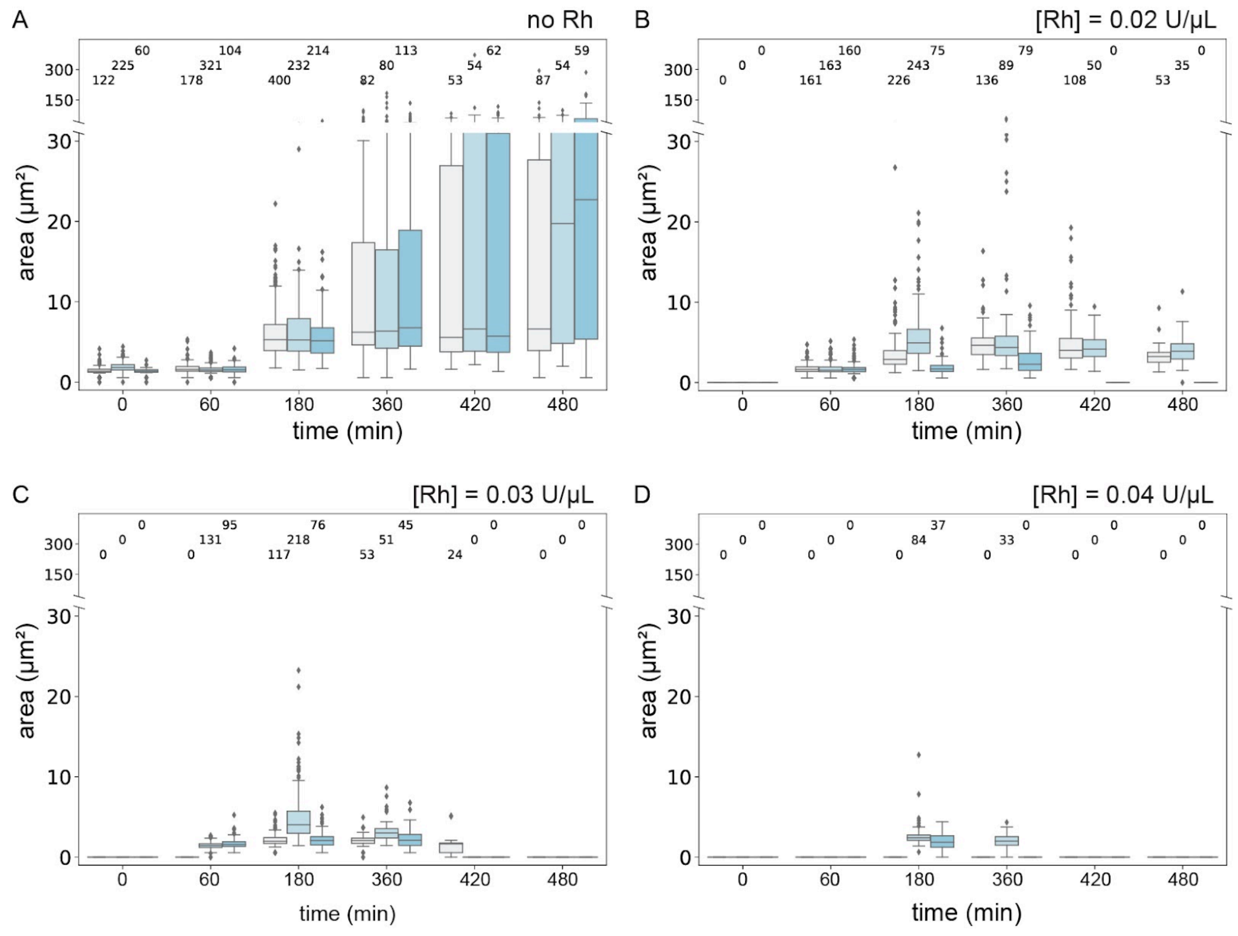


***Figure S12. Growth kinetic of DNA-RNA 4-arm nanostars in the presence of RNA-production and -degradation enzymes: box plots of the condensate size distribution for experiments reported in Figure 5 of the manuscript.*** *Images were taken 15, 60, and 180 minutes after addition of RNA-production (using RNAP) and -degradation (using RNase H) components. Box plots for the: experiments using design (D) that includes* $D_1$*,* $D_2$*,* $D_3$*, and* $R_4^{apt}$ *in the presence of SA and in the absence of RNase H; B) experiments using design (D) in the presence of both SA and 0.02 U/µL of RNase H. C) experiments using design (D) in the presence of both SA and 0.03 U/µL of RNase H; D) experiments using design (D) in the presence of both SA and 0.04 U/µL of RNase H. The distributions pool the data of triplicate, and the number of captured condensates corresponding to each time point is annotated inside the plots.*

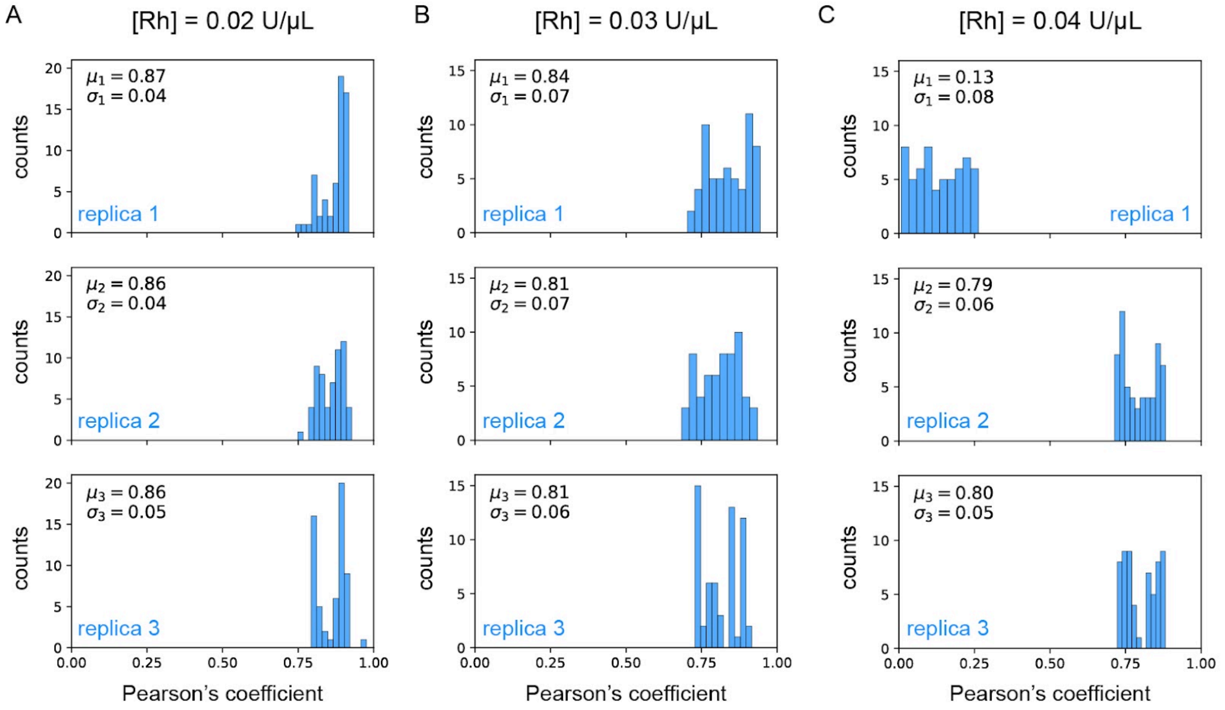


***Figure S13. Analysis of nanostars and protein colocalization 180 minutes after the coexistence of both RNAP and RNAse H enzymes.*** *Statistical distribution of Pearson's coefficient (r), for triplicate, of condensates 180 minutes after cotranscription initiation in the presence of RNAse H at: A) 0.02 U/µL; B) 0.03 U/µL; C) 0.04 U/µL. The coefficients close to 0 (A) indicate low colocalization of the FAM and Alexa 647 fluorophores (nanostar and protein, respectively), and consequently no protein recruitment. Contrarily, the r values close to 1 (B) indicate high colocalization, showing remarkable protein recruitment. The mean and standard deviation of r (µ and σ) for each replica are calculated from processing a random selection of 60 condensates.*

### References

(1) Agarwal, S.; Dizani, M.; Osmanovic, D.; Franco, E. Light-Controlled Growth of DNA Organelles in Synthetic Cells. *Interface Focus* **2023**, *13* (5), 20230017.
(2) Agarwal, S.; Osmanovic, D.; Dizani, M.; Klocke, M. A.; Franco, E. Dynamic Control of DNA Condensation. *Nat. Commun.* **2024**, *15* (1), 1915.
(3) Sato, Y.; Sakamoto, T.; Takinoue, M. Sequence-Based Engineering of Dynamic Functions of Micrometer-Sized DNA Droplets. *Sci Adv* **2020**, *6* (23), eaba3471.
(4) Fornace, M. E.; Huang, J.; Newman, C. T.; Porubsky, N. J.; Pierce, M. B.; Pierce, N. A. NUPACK: Analysis and Design of Nucleic Acid Structures, Devices, and Systems. **2022**. https://doi.org/10.26434/chemrxiv-2022-xv98l.
(5) Zadeh, J. N.; Steenberg, C. D.; Bois, J. S.; Wolfe, B. R.; Pierce, M. B.; Khan, A. R.; Dirks, R. M.; Pierce, N. A. NUPACK: Analysis and Design of Nucleic Acid Systems. *J. Comput. Chem.* **2011**, *32* (1), 170–173.
(6) Stewart, J. M.; Subramanian, H. K. K.; Franco, E. Self-Assembly of Multi-Stranded RNA Motifs into Lattices and Tubular Structures. *Nucleic Acids Res.* **2017**, *45* (9), 5449–5457.
